# Learning cognitive maps as structured graphs for vicarious evaluation

**DOI:** 10.1101/864421

**Authors:** Rajeev V. Rikhye, Nishad Gothoskar, J. Swaroop Guntupalli, Antoine Dedieu, Miguel Lázaro-Gredilla, Dileep George

## Abstract

Cognitive maps are mental representations of spatial and conceptual relationships in an environment. These maps are critical for flexible behavior as they permit us to navigate vicariously, but their underlying representation learning mechanisms are still unknown. To form these abstract maps, hippocampus has to learn to separate or merge aliased observations appropriately in different contexts in a manner that enables generalization, efficient planning, and handling of uncertainty. Here we introduce a specific higher-order graph structure – clone-structured cognitive graph (CSCG) – which forms different clones of an observation for different contexts as a representation that addresses these problems. CSCGs can be learned efficiently using a novel probabilistic sequence model that is inherently robust to uncertainty. We show that CSCGs can explain a variety cognitive map phenomena such as discovering spatial relations from an aliased sensory stream, transitive inference between disjoint episodes of experiences, formation of transferable structural knowledge, and shortcut-finding in novel environments. By learning different clones for different contexts, CSCGs explain the emergence of splitter cells and route-specific encoding of place cells observed in maze navigation, and event-specific graded representations observed in lap-running experiments. Moreover, learning and inference dynamics of CSCGs offer a coherent explanation for a variety of place cell remapping phenomena. By lifting the aliased observations into a hidden space, CSCGs reveal latent modularity that is then used for hierarchical abstraction and planning. Altogether, learning and inference using a CSCG provides a simple unifying framework for understanding hippocampal function, and could be a pathway for forming relational abstractions in artificial intelligence.

## Introduction

Vicarious trial and error [1], the ability to evaluate futures by mental time travel, is a hallmark of intelligence. To do this, agents need to learn mental models, or ‘cognitive maps’ [2, 3], from a stream of sensory information as they experience the environment around them [4]. Learning these mental abstractions is complicated by the fact that sensory observation is often aliased. Depending on context, identical events could have different interpretations and dissimilar events could mean the same thing [5]. As such, a computational theory for cognitive maps should: (1) propose mechanisms for how context and location specific representations emerge from aliased sensory or cognitive events, and (2) should describe how the representational structure enables consolidation, knowledge transfer, and flexible and hierarchical planning. Most attempts at developing such a theory, which include modeling hippocampus as a memory index, a relational memory space, a rapid event memorizer, and systems-level models of pattern-separation and pattern completion, have not reconciled the diverse functional attributes [6–8] of the hippocampus under a common framework. Recent models have attempted to reconcile the representational properties of place cells and grid cells using successor representation theory [9–11] and by assuming that these cells are an efficient representation of a graph [12]. Unfortunately, both these models fall short in describing how flexible planning can take place after learning the environment and are unable to explain several key experimental observations such as place cell remapping in spatial and non-spatial environments [13, 14] and the fact that some place cells encode routes towards goals [15, 16] while others encode goal values [17, 18].

A behaving agent often encounters external situations that look instantaneously similar, but require different action policies based on the context. In these situations, sensory observations should be contextualized into different states. In other times, dissimilar looking sensory observations might need to be merged on to the same state because those contexts all lead to same outcome. In general, to form a flexible model of the world from sequential observations the agent needs to have a representational structure and a learning algorithm that allows for elastic splitting and merging of contexts as appropriate [5, 19]. Moreover the representational structure should be such that it allows for dynamic planning, and handling of uncertainty.

Here we propose a specific higher-order graph – clone-structured cognitive graph (CSCG) – that maps observations on to different ‘clones’ of that observation as a representational structure that addresses these problems. We demonstrate that this structure can be represented as an extension of a probabilistic sequence model, and learned efficiently. CSCGs can explain a variety cognitive map phenomena such as discovering spatial relations from an aliased sensory stream, transitive inference between disjoint episodes of experiences, transferable structural knowledge, and shortcut-finding in novel environments. CSCG’s ability to create different clones for different contexts explains the emergence of splitter cells [15], and route-specific encoding [20], which we demonstrate using a variety of experimental settings common in neurophysiology. In a repeated lap-running task [21], CSCGs learn lap-specific neurons, and exhibit event-specific responses robust to maze perturbations, similar to to neurophysiological observations. CSCGs can also learn to separate multiple environments that share observations, and then retrieve them based on contextual similarity. Notably, the dynamics of clone-structure learning and inference gives a coherent explanation for the different activity remapping phenomena observed when rats move from one environment to another. By lifting the aliased observations into a hidden space, CSCGs reveal latent modularity that is then used for hierarchical abstraction and planning.

## Clone-structured cognitive graphs as a model of cognitive maps

The central idea behind CSCGs is dynamic Markov coding [22], which is a method for representing higher-order sequences by splitting, or cloning, observed states. For example, a first order Markov chain representing the sequence of events *A* − *C* − *E* and *B* − *C* − *D* will assign high probability to the sequence *A* − *C* − *D* (**Fig. 1a**). In contrast, dynamic Markov coding makes a higher-order model by splitting the state representing event *C* into multiple copies, one for each incoming connection, and further specializes their outgoing connections through learning. This state cloning mechanism permits a sparse representation of higher-order dependencies, and has been discovered in various domains [22–25]. With cloning, the same bottom-up sensory input is represented by a multitude of states that are copies of each other in their selectivity for the sensory input, but specialized for specific temporal contexts, enabling the efficient storage of a large number of higher-order and stochastic sequences without destructive interference. However, learning dynamic Markov coding is challenging because cloning relies on a greedy heuristic that results in severe suboptimality — sequences that are interspersed with zerothorder or first-order segments will result in an uncontrolled growth of the cloned states. Although [25] incorporated the cloning idea in a biological learning rule, the lack of a probabilistic model and a coherent global loss function hampered its ability to discover higher-order sequences, and flexibly represent contexts. An effective learning approach should split clones to discover higher order states, and flexibly merge them when that helps generalization.

**Fig. 1:**
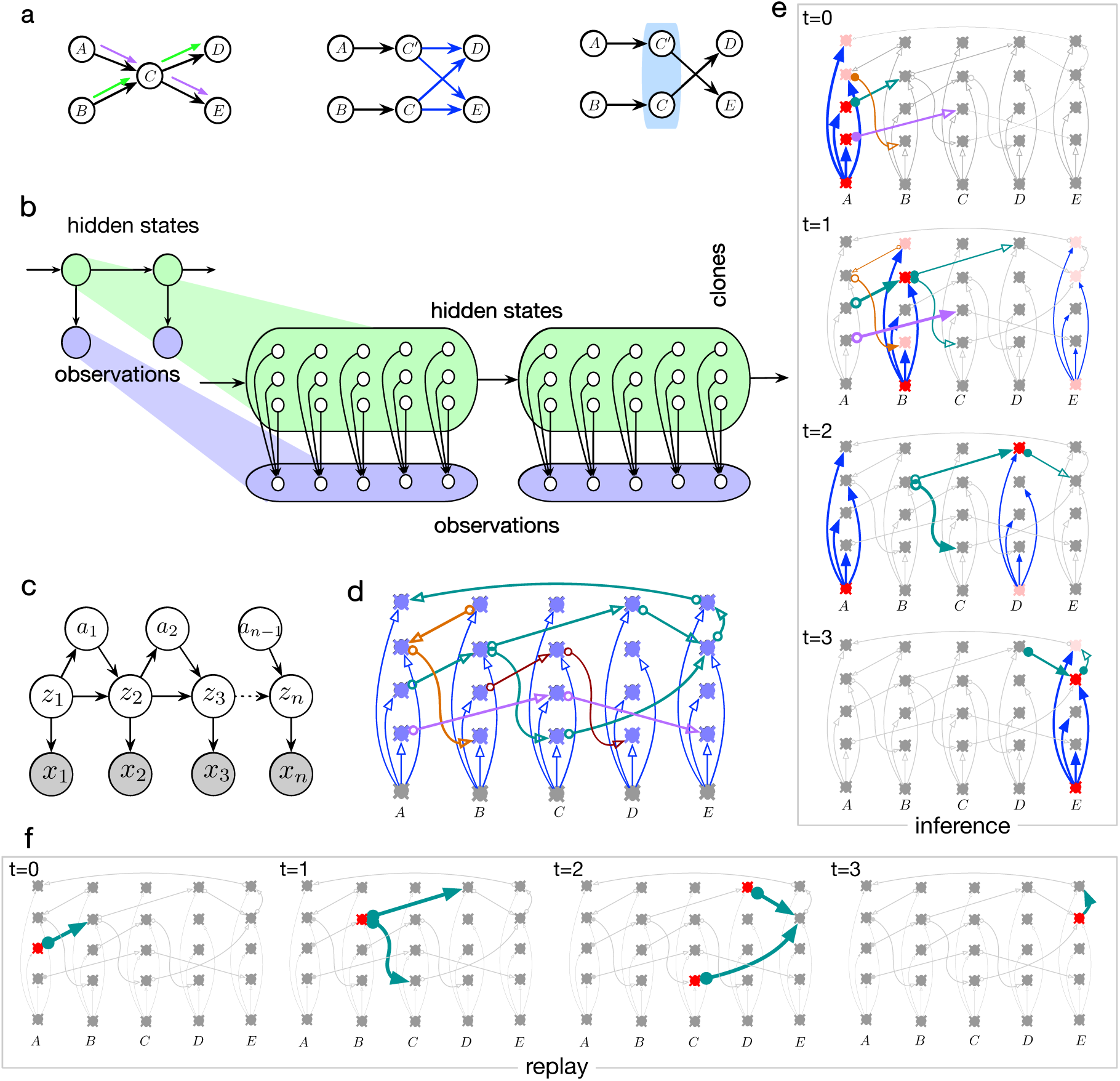
Clone-structured cognitive graph. **(a)** Sketch explaining dynamic Markov coding. A first order Markov chain modeling sequences A-C-E and B-C-D will also assign high probability to the sequence A-C-D. Higher-order information can be recovered by cloning the state C for different contexts. **(b)** Cloning structure of dynamic Markov coding can be represented in an HMM with a structured emission matrix, the cloned HMM. **(c)** CSCG extends cloned HMMs by including actions. **(d)** Neural implementation of cloned HMM. Arrows are axons, and the lateral connections implement the cloned HMM transition matrix. Neurons in a column are clones of each other that receive the bottom-up input from the same observation. **(e)** Inference dynamics in the cloned HMM neural circuit. Activations that propagate forward are the ones that have contextual (lateral) and observational (bottom-up) support. **(f)** Replay within the cloned HMM circuit.

Our previous work [26] showed that many of the training shortcomings of dynamic Markov coding can be overcome through cloned hidden Markov models – a sparse restriction of an overcomplete hidden Markov model (HMM) [27]. In cloned HMMs, the maximum number of clones per state is allocated upfront, which enforces a capacity bottleneck. Learning using the expectation-maximization (EM) algorithm figures out how to use this capacity appropriately to split or merge different contexts for efficient use of the clones to represent different contexts. In addition, cloned HMMs represent the cloning mechanism of dynamic Markov coding in a rigorous probabilistic framework that handles noise and uncertainty during learning and inference.

Both HMMs and cloned HMMs assume the observed data is generated from a hidden process that obeys the Markovian property. That is, the conditional probability distribution of future states, given the present state and all past states, depends only upon the present state and not on any past states. For HMMs, the joint distribution over the observed and hidden states given by the following equation:

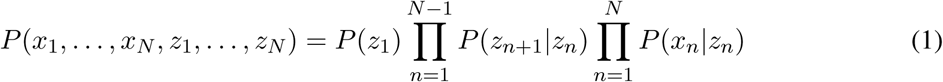

where *P* (*z*_1_) is the initial hidden state distribution, *P* (*z*_*n*+1_|*z*_*n*_) is the probability of transitioning from hidden state *z*_*n*_ to *z*_*n*+1_, and *P* (*x*_*n*_|*z*_*n*_) is the probability that observation *x*_*n*_ is generated from the hidden state *z*_*n*_. We assume there are *E* distinct observations and *H* distinct hidden states i.e. *x*_*n*_ can take a value from 1, 2, *…, E* and *z*_*n*_ can take a value from 1, 2, *…, H*.

In contrast to HMMs, in the cloned HMMs, many hidden states map deterministically to the same observation (**Fig. 1b**). The set of hidden states that map to a given observation are referred to as the *clones* of that observation. We use *C*(*j*) to refer to the set of clones of observation *j*. The probability of a sequence in a cloned HMM is obtained by marginalizing over the hidden states as follows:

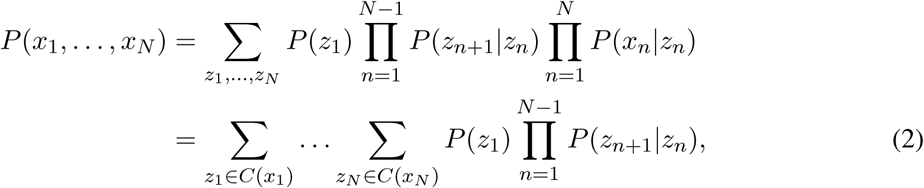

where the simplification is a result of *P* (*x*_*n*_ = *j*|*z*_*n*_ = *i*) = 0 for all *i* ∉ *C*(*j*) (and 1 otherwise). Moreover, since each hidden state is associated with a single observation, EM-based learning is significantly more efficient in cloned HMMs, allowing it to handle very large state spaces compared to standard HMMs [26]. See **Methods** for more details.

A hallmark of our model is the ability to handle noise and uncertainty via message-passing inference [28], and smoothing. Notably, just a forward and backward sweep of messages through the transition matrix *P* (*z*_*n*+1_|*z*_*n*_) is adequate for exact inference, and uncertainty about observations is handled through ‘soft-evidence’ messages. Smoothing [29] is a mechanism for incorporating robustness to noise and limited data in probabilistic models. In cloned HMMs, smoothing is accomplished by adding very small probability to some transitions that were unobserved in training. See **Methods** for more details.

### Neurobiological circuit

Like HMMs [30], cloned HMM can be readily instantiated as a neuronal circuit whose mechanistic interpretation provides additional insights on the advantages of the cloned representation. Each clone corresponds to a neuron, and the ‘lateral’ connections between these neurons form the cloned HMM transition matrix *P* (*z*_*n*+1_|*z*_*n*_). For example, the circuit in **Fig. 1d** shows how neurons can be connected in the cloned HMM to represent the following stored sequences *A* → *B* → (*C, D*) → *E* → *A* (green), *B* → *A* → *B* (light brown), *B* → *C* → *D* (dark brown), and *A* → *C* → *E* (purple).

The transition matrix can also be treated as a directed graph, with the neurons forming the nodes of the graph and the axonal branches forming the directed edges. The set of neurons that are clones of each other receive the same ‘bottom-up’ input (blue arrows) from the observation. The output of a clone-neuron is a weighted sum of its lateral inputs, multiplied by the bottom-up input, corresponding to the forward-pass message in HMM inference [30].

The evidence at any particular time instant can be uncertain (‘soft evidence’), manifesting as graded activation over the population of observation neurons. For a particular observation, the direct bottom-up connections from the observation to all its clones activate the the different sequences that observation is part of, and these activations are then modulated based on the specific contextual support each clone receives on its lateral connections. The population of clone neurons represent the probability of different contexts that are active at any time in proportion to their probability. **Fig. 1e** shows how these activities propagate for a noisy input sequence *A* → (*B, E*) → (*A, D*) → *E* from *t* = 0 to *t* = 3 corresponding to a true sequence *A* → *B* → *D* → *E*. The activations are represented in different shades of red, with lighter shades indicating weaker activations. At every time instant, the activated lateral inputs are highlighted, and these correspond to the clones active in the previous time step. By correctly integrating the context and noisy input, the clone activations of the cloned HMM filter out the noise to represent the true input sequence. **Fig. 1f** shows how sequences can be ‘replayed’ (sampled) from the circuit.

Queries like marginal or MAP inference can be implemented in neural circuits as forward and backward sweeps similar to the visualizations in Fig 1, analogous to the neural implementation of message-passing inference explored in earlier works [28, 30, 31]. The EM algorithm used for learning is well approximated by the neurobiological mechanism of spike-timing-dependent-plasticity (STDP) [32].

### CSCG: Action-augmented cloned HMM

CSCG extends cloned HMMs to include actions of an agent. An agent’s experience is a stream of sensation-action pairs (*x*_1_, *a*_1_), (*x*_2_, *a*_2_) *…* (*x*_*N*−1_, *a*_*N*−1_), (*x*_*N*_, −) where *x*_*n*_ ∈ Z^*^ are the agent’s sensory observations and *a*_*n*_ ∈ ℤ^*^ are the actions reported by the agent’s proprioception.

The observed actions are simply non-negative integers with unknown semantics (i.e., the agent observes *a*_1_ = 0 happened, but does not know that the action means ‘move north in the room’). In CSCG, the action is a function of the current hidden state and the future hidden state is a function of both the current hidden state and the action taken. The graphical model for this CSCG is depicted in **Fig. 1d**.

Mathematically, the joint observation-action density is:

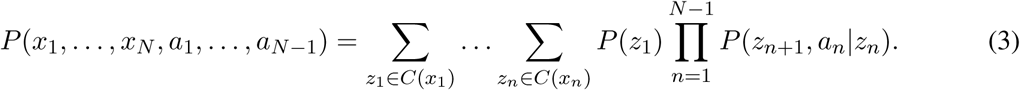

Our action-augmented model allows for the agent to learn which actions are feasible in a given state, compared to action-conditioned formulations [33] that only predict future observations from actions.

### Planning within a CSCG

Planning is treated as inference [34] and achieved using biologically plausible message-passing algorithms [28]. The goal can be specified as either a desired observation or as a specific clone of that observation. Planning is then accomplished by clamping the current clone and the target, and inferring the intermediate sequence of observations and actions required to reach these observations. It is easy to determine how far into the future we have to set our goal by running a forward pass through the graphical model and determining the feasibility of the goal at each step. The backward pass will then return the required sequence of actions. Importantly, because the graphical model is inherently probabilistic, it can handle noisy observations and actions with uncertain outcomes.

## Results

We performed several experiments to test the ability of CSCGs to model cognitive maps. We specifically tested for known functional characteristics such as learning spatial maps from random walks under aliased and disjoint sensory experiences, transferable structural knowledge, finding shortcuts, and supporting hierarchical planning and physiological findings such as remapping of place cells, and routespecific encoding.

### Emergence of spatial maps from aliased sequential observations

From purely sequential random-walk observations that do not uniquely identify locations in space, CSCGs can learn the underlying spatial map, a capability that is similar to people and animals. **Fig. 2a** shows a 2D room with the sensory observations associated with each location. The room has 48 unique locations, but only 4 unique sensory inputs (represented as colors), and an agent taking a random walk observes a sequence of these sensory inputs. A first-order sequence model would severely under-fit, and pure memorization of sequences will not learn the structure of the room because the same sequence hardly ever repeats. In contrast, a CSCG discovered the underlying 2D graph of the room perfectly (**Fig. 2b**). As the number of unique randomly placed observations increases, learning becomes easier (see Supplementary Results).

**Fig. 2:**
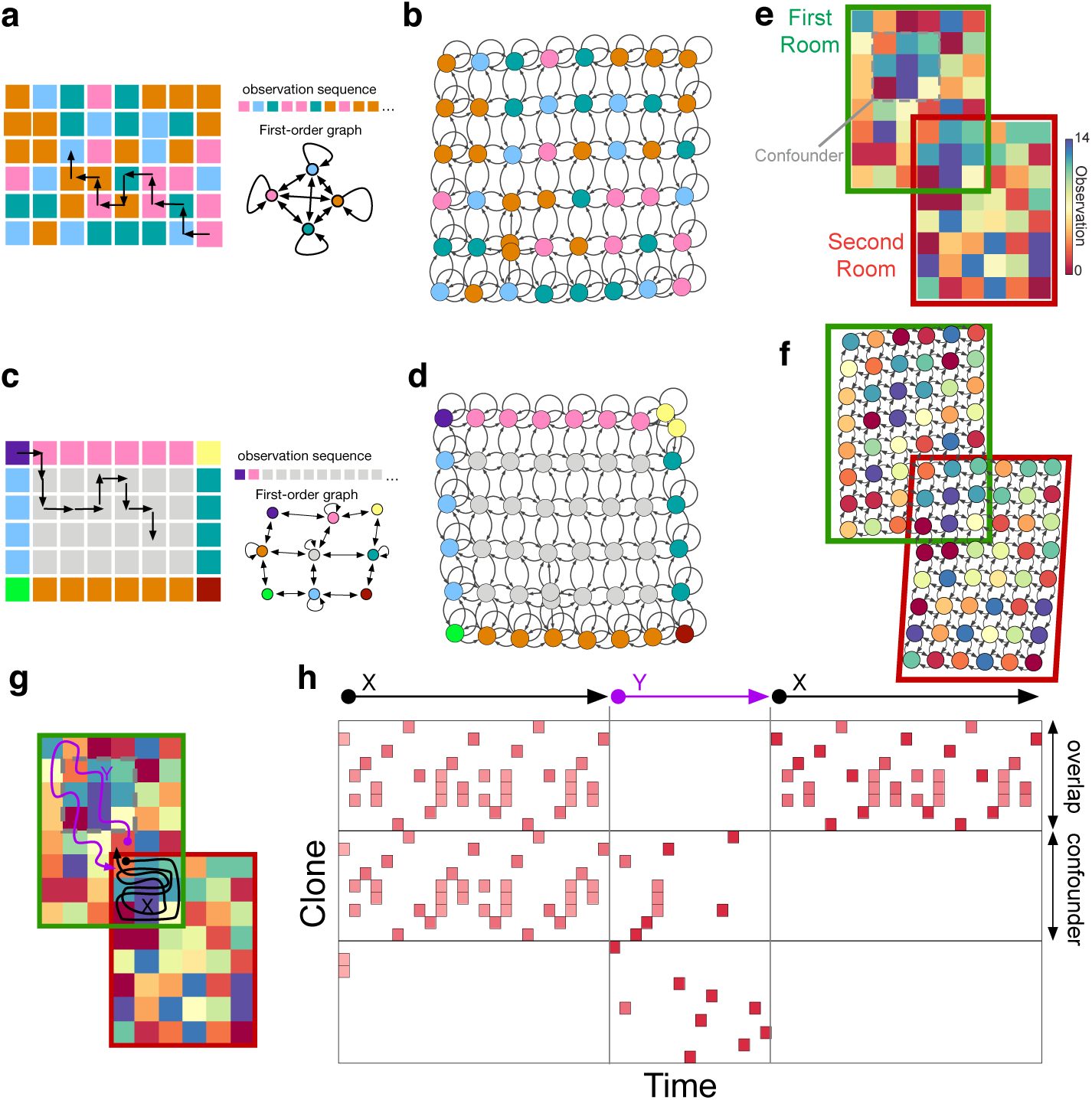
Spatial representations emerge from aliased sequential random-walk observations without Euclidean assumptions. **(a)** A random walk a room with only four unique observations (colors) will produce a sequence which is severely aliased as reflected in the first-order Markov chain. **(b)** In contrast, transition graph learned by CSCG on random walks in **(a)** recovers the spatial layout. Nodes in this graph are the clones, and the observation they connect to are indicated by the color of the node. **(c)** Room with a uniform interior produces aliased sequences highly correlated in time. **(d)** Transition graph learned by CSCG on random walks in **(c)**. Nodes are clones, and their observations are indicated by the node color. **(e)** An agent experiences two different, but overlapping rooms in disjoint sequential episodes. The overlap region also repeats in the first room, acting as a confounder. **(f)** As reflected in the transition graph, CSCG performs transitive inference to stitch together the disjoint experience into a coherent global map, and correctly positions the confounder. **(g-h)** Activation of clones over time as the agent takes the trajectories X (black), Y (purple), and X again in the maze shown in **(g)**. During the first traversal of X, the clones corresponding to the overlap and the confounding patch are active because the agent started within the overlap and stayed within. Stepping out of the overlap immediately resolves ambiguity, which is reflected in the clone activity during the traversal Y, and also during the second traversal of X. See also Supplementary Video 1.

Remarkably, CSCGs learn the spatial topology even when most of the observations are aliased like those from a large empty room where distinct observations are produced only near the walls as shown in **Fig. 2c**. The combination of high correlation between observations, and severe aliasing makes this a challenging learning problem. Despite this, the CSCG is able to perfectly learn the topology of the 6 × 8 room (**Fig. 2d**). This capability degrades as the room gets larger, but the degradation is graceful. For example, the periphery of a 9 × 11 room is well modeled, but the CSCG is unable to distinguish a few locations in the middle (see **Supplementary Results**).

### Transitive inference: disjoint experiences can be stitched together into a coherent whole

Transitive inference, the ability to infer the relationships between items or events that were not experienced at the same time, is attributed to cognitive maps [7]. Examples include realizing *A* > *C* from knowing *A* > *B* and *B* > *C*, or inferring a new way to navigate a city from landmarks and their relative positions experienced on different trips [35].

We tested CSCGs on a challenging problem designed to probe multiple aspects of transitive inference and found that it can stitch together disjoint episodes of sequential experience into a coherent whole. The experimental setting consisted of overlapping rooms (**Fig. 2e**), each with aliased observations like in the previous experiment. Moreover, the first room had an additional portion which was identical to the overlapping section between the two rooms. This design allows to test whether an agent that experiences only first room or second room exclusively and sequentially, can correctly figure out the relationship between the rooms and their overlaps. The combination of a large state-space, aliased observations, nested relationships, and two-dimensional transitivity makes the problem setting significantly harder than previous attempts [36]. We collected two independent sequences of action-observation pairs on each room by performing two separate random walks, and trained a single CSCG on both sequences. The result of training is visualized in **Fig. 2f**. The learned transition matrix (shown as a graph) has stitched together the compatible region of both rooms, creating a single, larger spatial map that is consistent with both sequences while reusing clones when possible. The confounding additional patch in the first room remains correctly unmerged, and in the right relative position in the first room, despite looking identical to the overlapping region.

Discovering the correct latent global map enables CSCG to make transitive generalizations. Although the agent has never experienced a path taking it from regions that are exclusive to Room 1 to regions exclusive to Room 2, it can use the learned map to vicariously navigate between any two positions in the combined space. Just like in the earlier experiment, the learning is purely relational: no assumptions about Euclidean geometry or 2D or 3D maps are made in the model.

Interestingly, plotting the activation of clones over time reveals that when the agent first traverses the overlapping region (trajectory *X* in **Fig. 2g**), clones corresponding to both the overlap region and the identical confounding region are active (**Fig. 2h**), indicating that the agent is uncertain of its position in the maze. This also suggests that the the agent’s belief in the cognitive map is split between the two possible realities (see **Supplementary Video 1**) because the overlap region and the confounding region are exactly the same without additional context. Stepping out of the overlap region gives the agent adequate context to resolve ambiguity. Subsequently, as the agent explores the confounding region (trajectory *Y* in **Fig. 2g**), clones corresponding to this region become more active, and the clones corresponding to the overlap region are no longer active. When the agent returns to the overlap region to follow the same sequence (trajectory *X*) it originally followed, the clone activities reflect that the agent is no longer confused between the overlap region and the confounding region.

### Learned graphs form a reusable structure to explore similar environments

The generic spatial structure learned in one room can be utilized as a schema [37] for exploring, planning, and finding shortcuts in a novel room, much like the capabilities of hippocampus based navigation [38]. To test this, we first trained the CSCG on Room 1 based on aliased observations from a random walk. As before, CSCG learned the graph of the room perfectly. Next, we placed the agent in Room 2 which is unfamiliar (**Fig. 3a**). We kept the transition matrix of the CSCG fixed, and re-initialized the emission matrix to random values. As the agent walks in the new room, the emission matrix is updated with the EM algorithm. Even without visiting all the locations in the new room, the CSCG is able to make shortcut travels between visited locations through locations that have never been visited (**Fig. 3b**). After a short traversal along the periphery as shown in **Fig. 3a**, we queried to find the shortest path from the end state to the start state. The CSCG returned the correct sequence of actions, even though it obviously cannot predict the observations along the path. Interestingly, Viterbi decoding [39] reveals the same hidden states that you would get if you Viterbi decoded the same path in Room 1. Querying the CSCG on the shortest path from the bottom left corner of the room to the start position, reveals the path indicated by the blue arrows in **Fig. 3b**. This solution is the Djikstra’s shortest path through the graph obtained from Room 1. Furthermore, if we ‘block’ the path we get another solution that is also optimal in terms of Djikstra’s algorithm (**Fig. 3c**). Even with partial knowledge of a novel room, an agent can vicariously evaluate the number and types of actions to be taken to reach a destination by reusing CSCG’s transition graph from a familiar room.

**Fig. 3:**
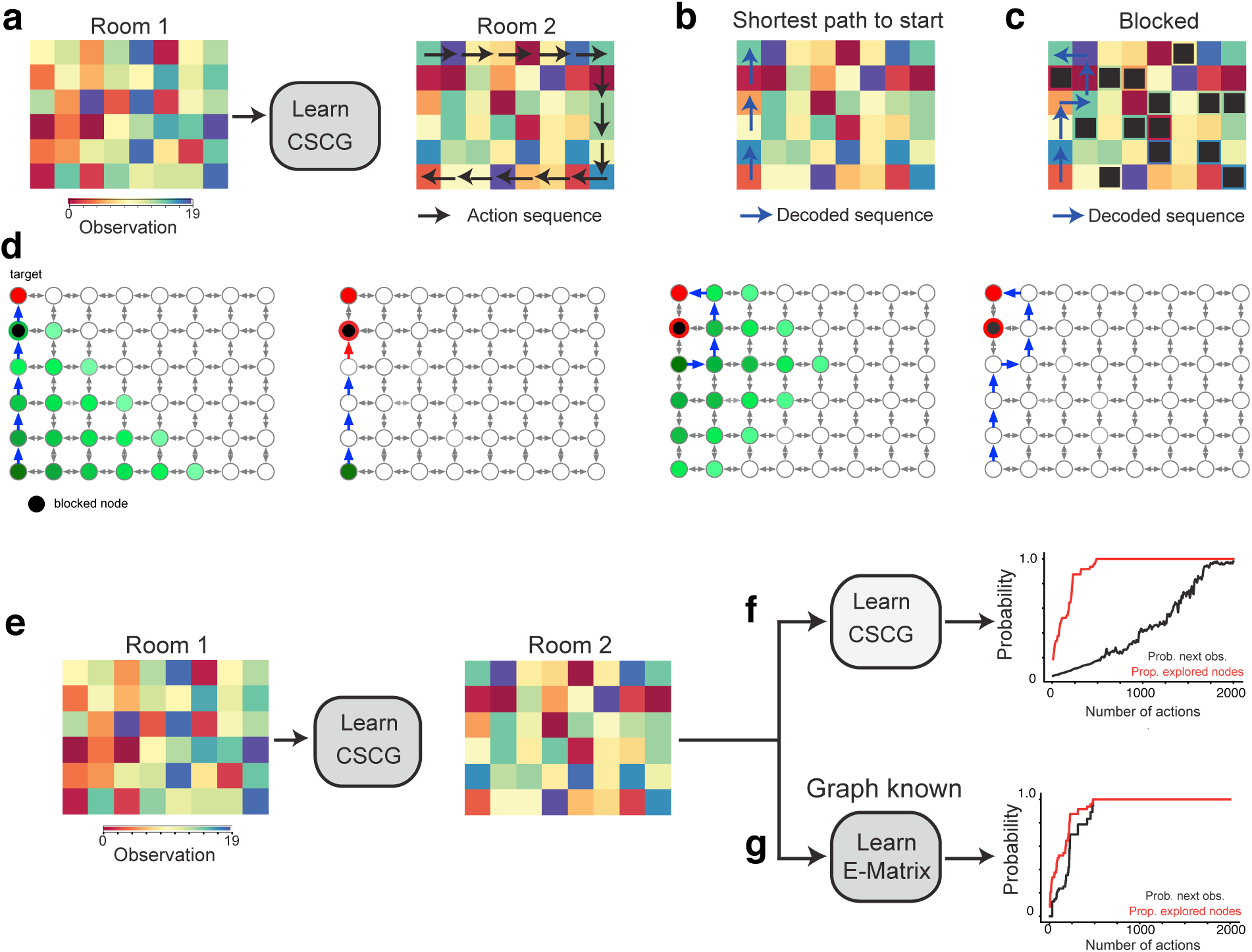
Learned transition graphs form a reusable schema to explore similar environments. **(a-c)** A CSCG trained on one room **(a)** and partial observations in a second, previously unseen room, utilizes the learned structure of the room to rapidly find both the shortest path to the origin **(b)** and navigate around obstacles **(c). (d)** Visualization of message propagation during planning and replanning. Messages propagate outward from the starting clone, and clones that recieve the message are indicated in green color. Lighter shades indicate messages that are later in time. The first plan is unaware of the obstacle, and the agent discovers the obstacle only when the action sequence is executed and a planned action fails (red arrow). This initiates a re-planning from the new location, and the new plan routes around the obstacle. **(e-f)** The transition matrix (graph) learned in one room can be used as a re-usable structure to quickly learn a new room with similar layout but different observations.

When the transition matrix from the old room is reused, the new room is learned very quickly even when the agent explores using a random walk: the new room is learned fully when all the locations in the room are visited at least once (**Fig. 3d-f**). The plots show the proportion of the room explored and the average accuracy of predicting the next symbol as a function of the number of random-walk steps.

### Representation of paths and temporal order

CSCGs learn paths and represent temporal order when the observed statistics demand it, for example when the observations correspond to an animal repeatedly traveling prototypical routes. For example, consider the T-maze shown in **Fig. 4a**, which is traversed in a *figure-of-eight* pattern either from the right (blue path) or the left (red path). As a result, the two paths share the same segment. Interestingly, CSCG learns separate clones for this shared segment (**Fig. 4b**) and similar to the observations in [15], the activity of clones in this overlapping segment will indicate whether the rat is going to turn left or right (**Fig. 4c**). It is important to note that the ability of CSCGs to learn flexible higher-order sequences is independent of the modality [4]. In particular, the inputs can correspond to spatial observations, odors, sequences of characters, or observations from any other phenomenon [26]. CSCG will learn an approximation of the graph underlying the generative process, in close correspondence with the role for cognitive maps envisaged by [2]. We illustrate in **Fig. 4e** the CSCG learned for a maze with a shared path shown in **Fig. 4d**.

**Fig. 4:**
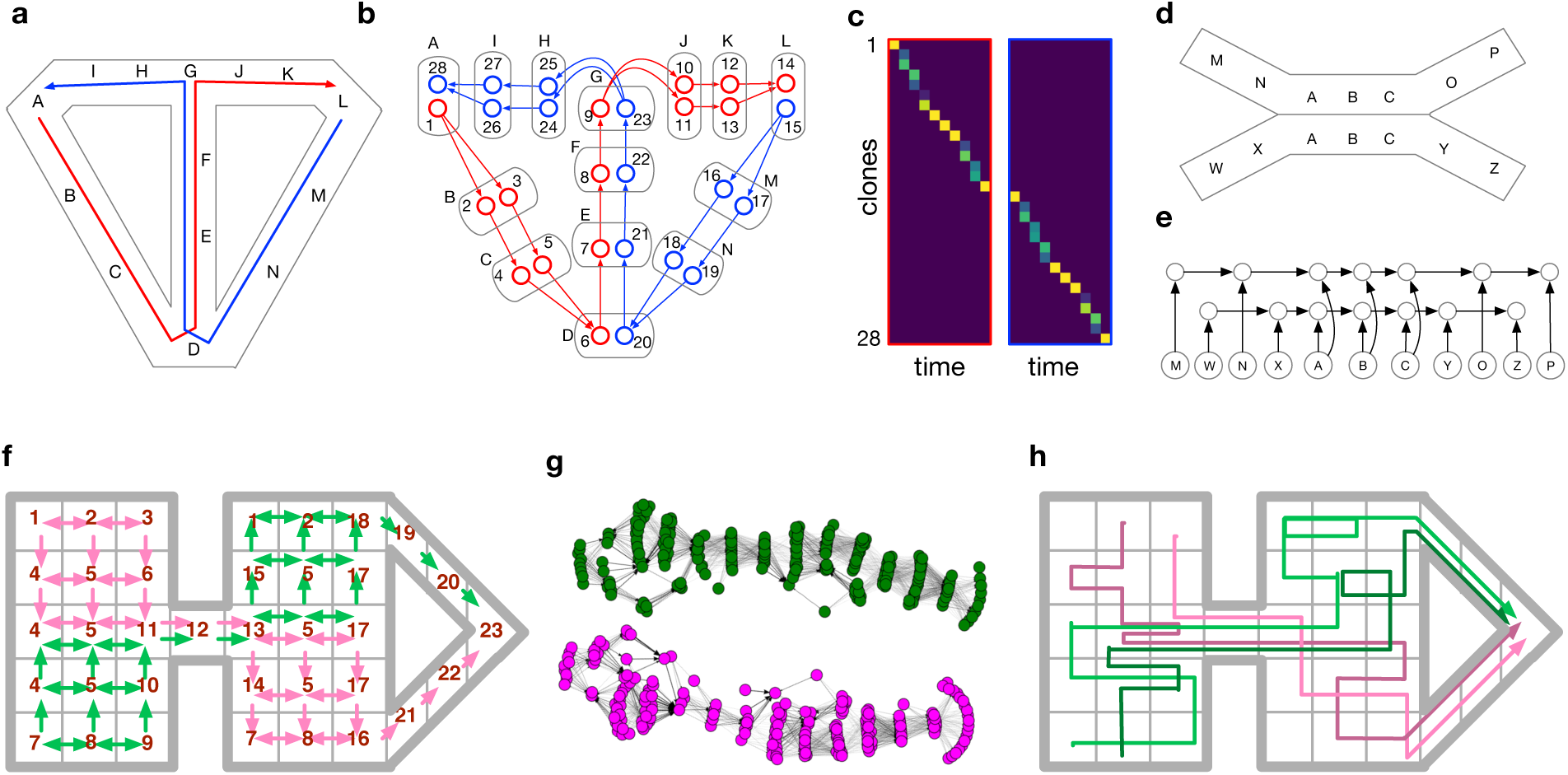
Learning temporal order and paths. In all experiments, CSCG learned the optimal model for prediction, and the learned circuits matched neurobiological observations. **(a)** Modified T maze from [15] with an overlapping segment between the blue and red paths. **(b)** CSCG learns separate clones for the two routes passing through the overlapping segment. Similar to the observations in [15], the activity of clones in this overlapping segment will indicate whether the rat is going to turn left or right. **(c)** Activity of the clones for the right trial, and the left trial. Distinct neurons are active in the overlapping segment for left-turn trials vs right-turn trials although the observations in the overlapping segment are identical for both trials. Note that clones are not limited to one time step. CSCG learning is able to propagate clones backward into multiple time steps to unravel long overlapping paths. **(d)** Overlapping odor sequences from [40] **(e)** Full circuit learned by the CSCG shows that it has learned distinct paths in the overlap, as in [40]. **f)** A complex maze in which the agent takes two stochastic paths indicated in magenta and green. Observations in the maze are marked by numbers and, as before, the same observation can be sensed in many parts of the maze. The green and magenta paths overlap in up to seven locations in the middle segment (observations 4-5-11-12-13-5-17). The stochasticity of the paths and the long overlaps make this a challenging learning problem. In contrast to mazes in **(a)** and **(d)**, the two paths in this maze lead to the same destination as in [20] **(g)** Transition graph learned by the CSCG shows that 2 different chains are learned for the 2 routes in **(f)**, similar to the observation that place cells encode routes, not destinations [20]. **(h)** Paths replayed from the CSCG after it was trained on sequences from **(f)**. As they pass through the overlapping segment, the green and magenta routes maintain the history of where they originated.

Neurophysiological experiments have shown the emergence of ‘splitter cells’ in the hippocampus [15]. These cells represent paths to a goal rather than physical locations and emerge as rats repeatedly traverse the same sequential routes as opposed to taking random walks [20]. **Fig. 4f** shows a maze in which the agent can traverse two different routes (indicated by the magenta and green lines) to reach the same destination. Both these routes have regions in which the exact path that the agent follows is stochastic, as denoted by the arrows that indicate the possible movements from each cell. Observations in the maze are marked by numbers and, as before, the same observation can be sensed in many parts of the maze. Additionally, the two routes intersect and share a common segment. CSCGs trained on these paths are able to represent both routes by using different clones for each of the routes, analogous to the route dependency exhibited by place cells in similar experiments. We observe that disjoint subsets of clones will activate when traversing each of the routes. **Fig. 4g** shows that when conditioning on the starting state, sampling in the learned CSCG will always produce paths that are consistent with the two routes. By visualizing the graph defined by the CSCG transition matrix, we see that the two routes are represented with two different chains (**Fig. 4g**). With a first-order model, when the shared segment is reached, all context about the previous segments will be lost and the model will make incorrect predictions about the future path. CSCGs, on the other hand, are able to capture the history of the path and therefore properly model the routes and their distinct start states.

Learning higher-order sequences in a CSCG can also explain recently discovered phenomena like chunking cells and event-specific representations (ESR) [21], place cell activations that signal a combination of the location and lap-number for different laps around the same maze. **Fig. 5a** shows a setting similar to the experiment in [21] where a rat runs four laps in a looping rectangular track before receiving a reward. A CSCG exposed to the same sequence learned to distinguish the laps and to predict the reward at the end of the 4th lap. Planning for achieving the reward recovered the correct sequence of actions, which we then executed to record the activations of the clones in different laps. Visualizing the propagation of beliefs of each clone, either conditioned on the observation or the action, produces a sequence-like activation pattern where one clone is active for each sensory observation, and as such the different laps around the maze are encoded by different clones (**Fig. 5b**). Similar the neurons in the hippocampus, whose firing rates are shown in **Fig. 5c** [21], clones show graded activity across laps. A clone is maximally active for an observation when it occurs in its specific lap, but shows weak activations when that observation is encountered in other laps, a signature of ESR. This occurs naturally in the CSCG due to smoothing and the dynamics of inference, visualized in **Fig. 5e**. Sun and colleagues reported that despite extending the maze, neurons in the hippocampus still respond uniquely to each lap. We mimicked this experiment by elongating our maze in one dimension, by introducing repeated, or aliased, sensory observations (**Fig. 5d**). Again, as with the smaller maze, we observed that clones were uniquely active on each lap and parsed each lap as a separate contextual event (**Fig. 5d**). In this particular example, the cognitive map for this maze is a chain of observations (see **Fig. 5e**) which split each lap into distinct contextual events. In doing so, the agent is able to identify which lap it is in based on identical local observations. Robustness of ESR to maze elongations can also be explained by inference in a smoothed CSCG – a repeated observation is explained as noise in the previous time step, and re-planning from the current observation recovers the correct sequence of actions.

**Fig. 5:**
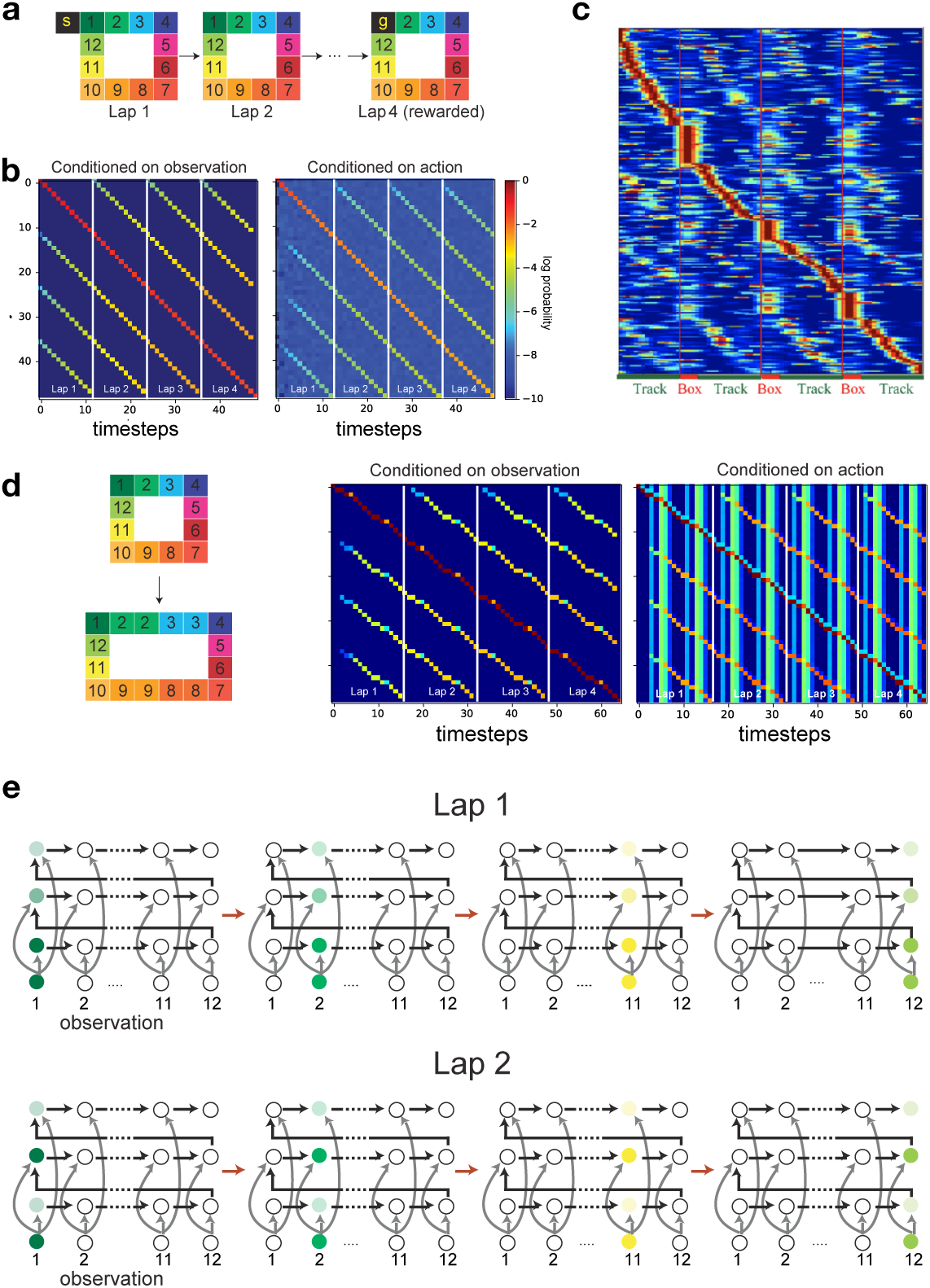
Lap-neurons and event-specific representations. **(a)** A CSCG was trained on observations from four laps around a square maze similar to [21]. It learned to predict the laps perfectly, including the reward at the end of the 4th lap, and planning to get the reward returned the correct sequence of actions. **(b)** Clone activations for the 4 different laps. Rows correspond to neurons (clones). The activations show that there are different clones that are maximally active for different laps, but the other clones are partially active at their corresponding locations, similar to the neurophysiological observations in [21] regarding event-specific-representations. **(c)** Place cell traces from [21], included with permission. **(d)** The event-specific representations persist even when the maze is elongated. The CSCG is not trained on the elongated maze. **(e)** Visualization of the circuit learned by the CSCG including the transition graph, connections from the observations, and activation sequences. The CSCG learned one clone per lap for each position. Smoothing in the CSCG explains why other clones of other laps are partially active. See text for details.

### Learning multiple maps and explaining remapping

Remapping is the phenomenon where hippocampal place cell activity reorganizes in response to a change in the physical environment. Remapping, which can either be global or partial [19, 41–44], depends on how the hippocampus can segregate, store, and retrieve maps for multiple environments that might be similar or dissimilar [13, 41].

Similar to the hippocampus [19], CSCGs can learn to separate multiple maps from highly similar environmental inputs, represent those maps simultaneously in memory, and then use contextual similarity to retrieve the appropriate map to drive behavior. In **Fig. 6a**, we show 5 different 5 × 5 rooms that all share the same 25 observations, but arranged differently in space. We learn a single CSCG from sequences of random walks in each of these mazes where the walks switched between different rooms at irregular intervals, without providing any supervision about the maze identity or time of switching.

**Fig. 6:**
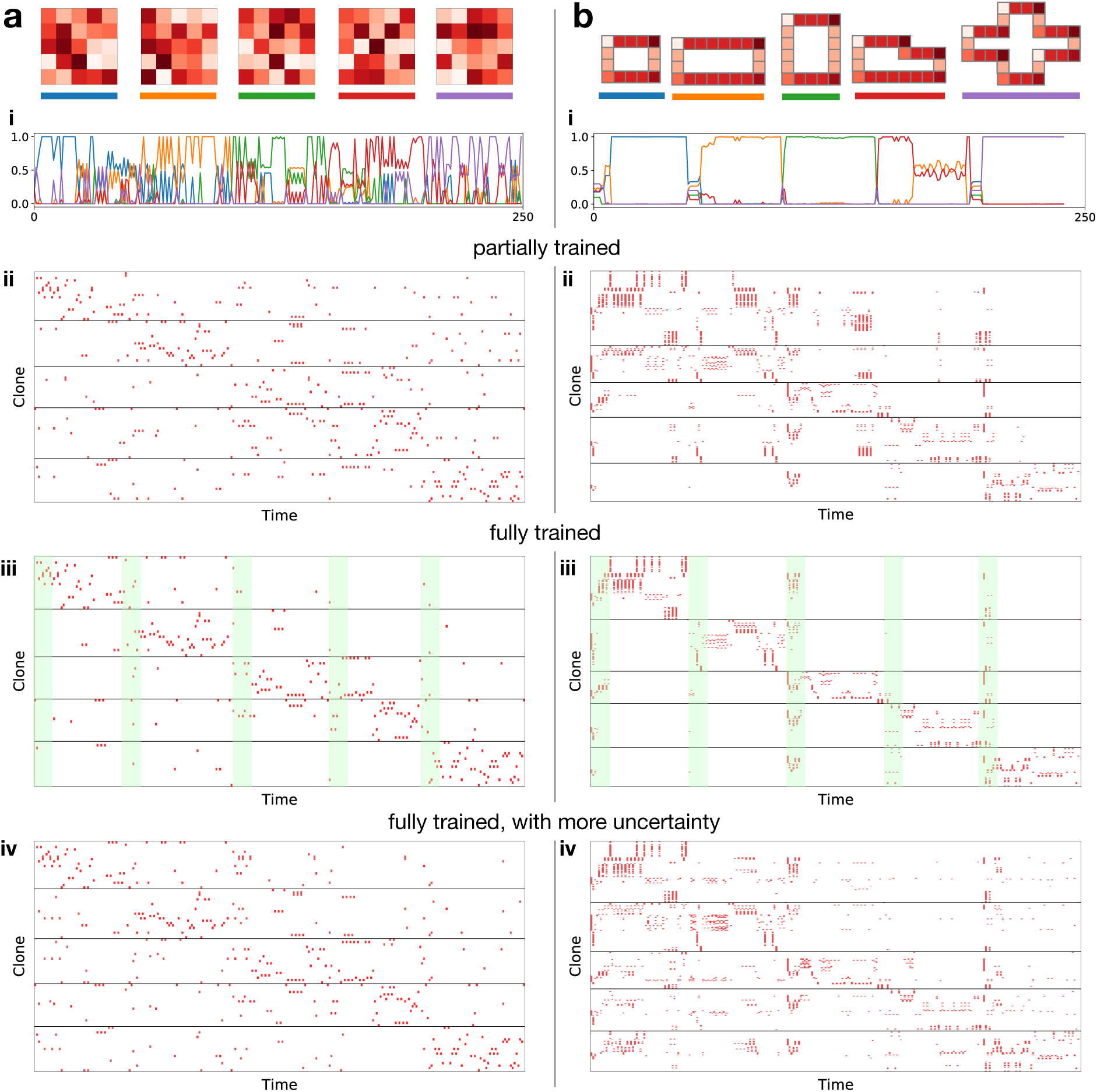
Remapping. Sets consisting of five different rooms **(a)** & mazes **(b)** mazes are used to study activity remapping. In set **(a)** the 5 rooms share 25 different observations arranged randomly, and in set **(b)** the 5 mazes share 6 observations arranged in geometrical shapes. Row **(i)**: inferred room/maze belief. Rows **(ii)-(iv)** Clone activity traces for a random walk of 50 steps each in rooms (mazes) 1 to 5 under different conditions (partially trained, fully trained, and more uncertainty). All traces are based on the same random walk and use the same clone ordering. Fully trained CSCG produces global remapping, and partially trained CSCG produces partial remapping. Adding more uncertainty to a fully trained CSCG produces rate remapping.

Although all observations are shared between the mazes, the CSCG learns to form different clones for the different rooms. Figure **Fig. 6ai** plots the agent’s belief about which map it is in as it goes through a 50-step random walk sequence in each room from the first to the last, showing that the maze identity is represented in the population response, despite the ambiguous instantaneous observations.

We conducted a series of experiments to evaluate how the similarity between mazes, predictability within each maze, the amount of learning, and the amount of noise and uncertainty affect the degree of reorganization of neural responses. These experiments used two sets of environments — mazes and rooms. Rooms are the 5×5 rooms described earlier (**Fig. 6a**), mazes consist of 5 different shapes (**Fig. 6b**) composed of 6 distinct observations (4 different corners, and vertical or horizontal arms). The mazes have better within-maze predictability compared to the rooms because of the lower-branching factor of the random walk, and mazes are more similar to each other compared to the similarity between different rooms. For each of these sets, we trained a CSCG, and evaluated how remapping changed with the amount of training, and uncertainty (see **Fig. 6a i-iv** and **Fig. 6b i-iv**).

Our results suggest that global remapping, partial remapping, and rate remapping can be explained using CSCGs: they are manifestations of learning and inference dynamics using a cloned structure when multiple maps are represented in the same model. We were able to reproduce different remapping effects by varying the amount of training and uncertainty. The rows (ii) to (iv) in **Fig. 6a-b** show the neural responses of two CSCGs that learned to represent the corresponding rooms and mazes. All the neural traces in a column correspond to the same random walk where the agent takes 50 steps in each room/maze, from the first to the last. When the CSCG is fully trained until the EM algorithm converges, the neural responses from the different mazes overlap the least, producing an effect similar to global remapping (**Fig. 6aiii and biii**) [41]. If the CSCGs are partially trained, the clones only partially separate – while many remain exclusive to particular mazes or rooms, a large number are also active in multiple mazes/rooms (**Fig. 6aii and bii**), corresponding to the effect of partial remapping [13, 42]. In a fully trained model, more smoothing, or soft-evidence that reflects uncertainty, creates neural responses similar to rate remapping [13, 44](**Fig. 6aiv and biv**): all the neurons that fire in the fully trained case still fire in this case, but with a lowered rate of firing. This occurs because uncertainty and smoothing causes more sharing of the evidence among clones that represent the same observation.

The similarity between the rooms (mazes), and the amount of predictability within each room (maze), also affects the dynamics of remapping. This can be observed by comparing the traces for the rooms with that of the mazes in **Fig. 6a, b**. In **Fig. 6bi**, the beliefs within each maze are more stable compared to those in the rooms due to the stricter temporal contexts in the mazes [19]. Fluid temporal contexts in rooms produce more progressive deformation of beliefs [45]. The structural similarity between the different mazes results in a longer transient period right after entering a new maze resulting in non-instantaneous switching of beliefs [45]. This is also reflected in **Fig. 6bii-iv**, where clones in multiple mazes are active at the point of switching (green bars).

Taken together, our experiments demonstrate the conditions and mechanisms that determine how the hippocampal network may abruptly switch between pre-established representations or progressively drift from one representation to the other, producing a variety of remapping effects.

### Community detection and hierarchical planning

Humans represent plans hierarchically [46]. Vicarious evaluations involve simulating paths to a goal, and hierarchical computations make these simulations tractable by reducing the search space [47]. To enable hierarchical planning, the learning mechanism should be able to recover the underlying hierarchy from sequentially observed data.

By learning a cloned transition graph, CSCG lifts observations into hidden space, enabling discovery of graph modularity that might not be apparent in the observation. Community detection algorithms [48], can then partition the graph to form hierarchical abstractions [6] useful for planning and inference. Like planning, and inference in CSCGs, community detection can also be implemented using message-passing algorithms [49] that make them biologically plausible [28].

We tested CSCGs for their ability to learn hierarchical graphs by simulating the movement of an agent in two mazes. The first maze is a modular graph with three communities where the observations are not unique to a node (**Fig. 7a**), in contrast to earlier studies using this graph [6, 9] where observations directly identified the nodes. Due the degeneracy of observations, community detection or MDS on the SR matrix fails to reveal the hidden communities (**Fig. 7b**). In contrast, community detection on CSCGs trained from random walks readily reveal the correct community structure. The second maze, shown in **Fig. 7d**, has a total of sixteen rooms arranged as a 4×4 grid. Each room has aliased observations, and are connected by corridors (black squares). The aliasing is global: instantaneous observations do not identify the room, corridor, or location within a room. Additionally, the maze is structured in such a way that there are four hyper-rooms making this maze a three-level hierarchy. As in the earlier examples, training a CSCG on random walk sequences learned a perfect model of the maze. We then used community detection to cluster the transition matrix of the CSCG (**Fig. 7e**). This clustering revealed a hierarchical grouping of the clones (**Fig. 7f**), and a connectivity graph between the discovered communities. The communities respected room boundaries: although some rooms were split into two or three communities, no community straddled rooms. Applying community detection once again on this graph revealed the four hyper-rooms (**Fig. 7f**) which were the highest level of the hierarchy. To navigate to a particular final destination *F* from a starting location *S* using this map, the agent first has to identify in which of these four rooms the goal is located, then plan a route in the community graph between the source community and the destination community (**Fig. 7h**). In doing so, the search-space in the lower level graph is significantly reduced, making planning in the hierarchical CSCG-learned graph more efficient than planning directly in the original graph. We implemented this form of hierarchical planning and found that we were always able to recover an efficient path between randomly selected start and end position (See Supplementary Methods for more details).

**Fig. 7:**
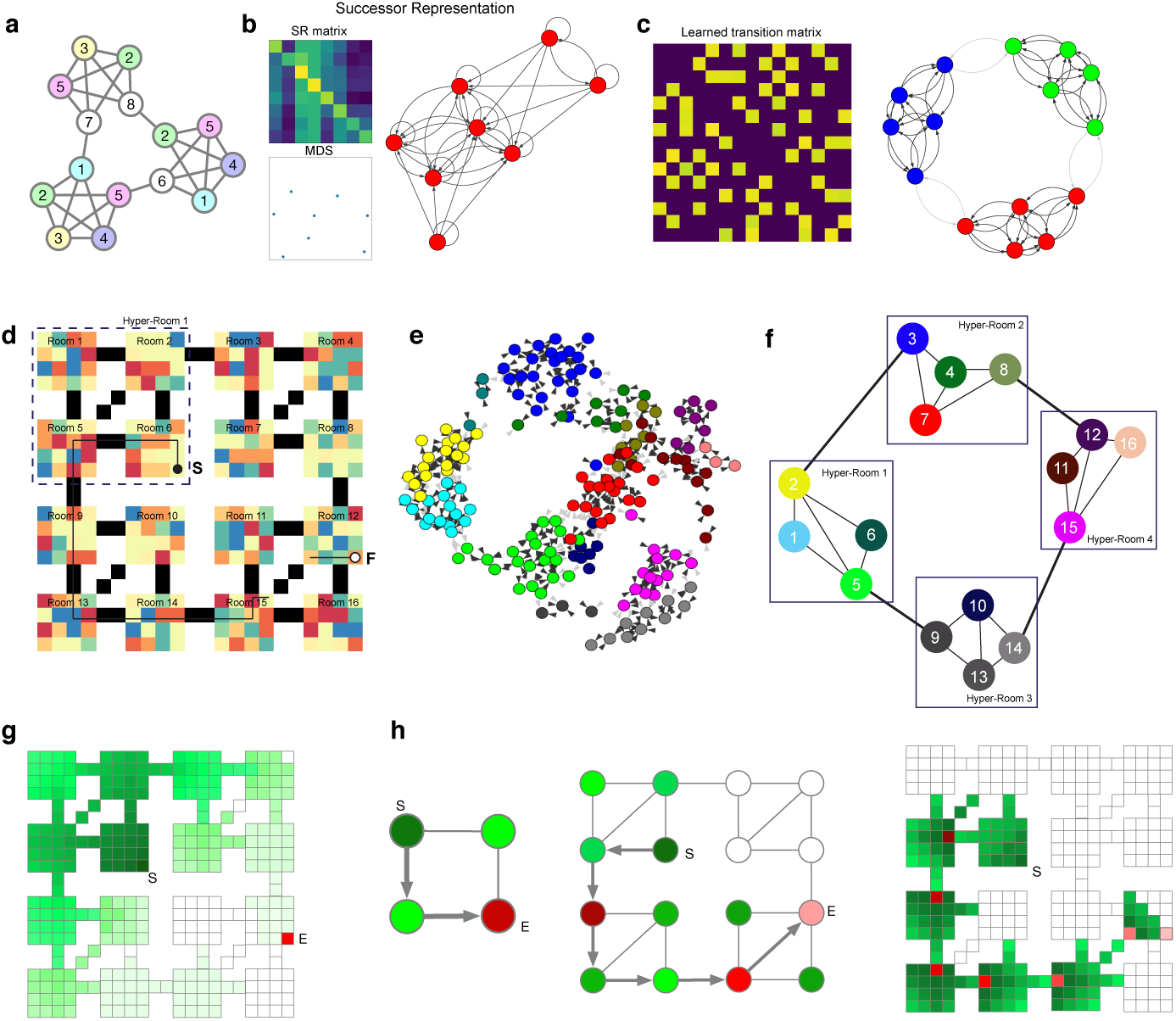
CSCGs enable hierarchical abstraction and planning. The cloned graph of the CSCG lifts the observations into a hidden space, allowing for discovery of modularity that is not apparent in the visible observations. **(a)** The modular graph from [6], modified to have aliased observations. Observations at each node are indicated by the numbers, and many different nodes produce the same observation. **(b)** MDS or community detection on the SR matrix of random walks in **(a)** does not reveal the modularity of the graph. **(c)** Community detection on the CSCG transition matrix successfully recovers the modularity of graph in **(a)**, recovering three communities. **(d)** A maze that has an embedded 3-level hierarchy. Sensory observations are aliased both within rooms and across rooms. The black pixels denote ‘bridges’ between the rooms. CSCG is trained on random walks from this maze. Community detection on the learned CSCG transition matrix revealed a first level of organization into rooms **(e)**, and another level of community detection revealed hyper rooms **(f)**, resulting in a 3-level hierarchical graph reflecting the nested structure of the maze. Planning a path (black arrows) between two rooms (denoted as (S)tart (filled black dot) and (F)inish (open black dot) in **(d)**) was achieved by finding the shortest path between hyper-rooms to navigate, next finding the shortest path between rooms, and lastly finding the shortest path within the rooms in this reduced search space. **(g)** Visualization of planning message propagation in the one-level graph. Messages propagate in the whole maze, indicating a wide search area. **(h)** Visualization of hierarchical planning. Routes are first identified on the highest level, which then becomes sub-goals at the lower level. The red colored nodes indicate the sequence of sub-goals, and their intensities reflect the ordering of the subgoals. Compared to **(g)**, hierarchical planning requires less messages to be propagated so it is faster.

Learning higher-order graphs that encode temporal contexts appropriately is crucial for the extraction of the hierarchy using community detection algorithms. Approaches that learn first-order connectivity on the observations, for example successor representations on observations [10], will not be able to form the right representations because the observations are typically severely aliased (see **Supplementary Fig. 3**).

## Discussion

Current theories of how cognitive maps are learned from sensory inputs and how they are used for planning have not been able to reconcile the vast body of experimental evidence. In this paper we pursued the strong hypothesis that hippocampus performs a singular algorithm that learns a sequential, relational, content-agnostic structure, and demonstrated evidence for its validity [4]. Through a series of experiments, we demonstrated how the CSCG is able to store, abstract, and access sequential sensory experience [4, 50]. Realizing this core idea required several interrelated advancements: (1) a learning mechanism to extract higher-order graphs from sequential observations, (2) a storage and representational structure that supports transitivity, (3) efficient context-sensitive and probabilistic retrieval, (4) and learning of hierarchies that support efficient planning – techniques we developed in this paper. As a model CSCG spans multiple levels of the Marr hierarchy. The computational specification is based on probabilistic models and optimal inference, and its algorithmic realization utilized neuroscience insights [24]. Moreover, the graphical model and the algorithmic realizations of its learning and inference readily translate to a neurobiological implementation that offers mechanistic explanations for all the experimental phenomena we considered.

CSCGs differ substantially from the Tolman-Eichenbaum Machine (TEM) [33, 51], a recent model on structure learning for hippocampal circuits. CSCGs can solve the tasks considered by TEM plus others as demonstrated in this work. For instance, unlike TEM, CSCGs can plan to achieve arbitrary goals selected at test time (see **Fig. 3b-c**) and natively handle erroneous or ambiguous observations (see **Retrieval and Remapping** in the supplementary material). CSCGs also allow for efficient exact inference, which enables sophisticated queries to be answered quickly and exactly. In contrast, the representational complexity of TEM only allows for approximate inference, and requires a higher computational effort. For instance, the problem in **Fig. 5a** uses 4 laps of 12 steps each, and is solved in seconds on a single CPU core, whereas for an equivalent problem to be solvable with TEM, it needs to be simplified to 3 laps of 4 steps each. CSCGs are natively probabilistic and handles uncertainty and noise, whereas current TEM realizations do not. Most importantly, the abilitiy of CSCGs to lift observations into a latent graph that reveals modularity offers it a powerful advantage over TEM by enabling the formation of abstraction hierarchies, see **Fig. 7**.

A commonly used theory for hippocampal function is the successor representation framework [9, 10, 52] which represents the current state of an agent by aggregating distributions over its future locations for a given policy. However, this places several limits on the representation. First, due to temporal aggregation, ordering in time is lost. Additionally, successor representations do not allow separate access to the current location and future locations, and mix the order of future locations [53]. In contrast, CSCGs provide separate access to the present and the predicted future and preserves ordering, a property vital for effective planning. Second, successor representations are a function of the policy. It is emphasized [9] that the value function can be easily recomputed when the reward changes, without recomputing the successor representation. However, what really needs to change when the reward changes is the policy, which in turn, requires the successor representation to the recomputed. Since CSCGs capture the dynamics of the world, they can update the policy on the fly. The observation of grid cell-like properties in the eigenvectors of the successor representations could be a property of all methods that employ a transition matrix (see **Supplementary Results**), and we suspect that this property in itself might not have any behavioral relevance. Finally, although the successor representation can be used to find communities, it requires the world to be fully observable without latent states. In contrast, CSCGs have the ability to split aliased observations into different contexts to discover latent graphs and communities.

CSCGs have intriguing connections to schema networks [54]. Like schema networks, CSCGs encode relational knowledge. Creating different clones for different temporal contexts is similar to the idea of *synthetic items* used to address state aliasing [55]. We intend to explore these connections in future work. Schema cells have been observed in the hippocampus [37], and CSCGs might be able to explain their emergence and properties. In addition, since sequence learning takes place in many other brain areas, for example the parietal cortex [56] and the orbito-frontal cortex [57], a natural extension of this work would involve learning higher order conceptual relationships and applying them to cognitive flexibility. The present work can be further extended by combining it with the active inference frame work [58] which provides a guiding principle for combining exploration and exploitation. Using active inference, at the beginning of learning, an agent will be driven by exploration because its world model is very uncertain, and will slowly increase the amount of exploitation as its knowledge of the world increases. Although active inference has been so far used in far simpler models that would not be able to solve the experiments presented in the current work, CSCG’s probabilistic formulation is compatible with representing the certainty of the model using hierarchical priors over model parameters, offering an avenue for future research.

In concordance with [50], CSCGs represent sequences of content-free pointers: each pointer can be referring to a conjunction of sensory events from different modalities. The output from grid cells, path integration signals, is treated as just another sensory modality. Grid cell outputs provide a periodic tiling of uniform space, which is advantageous for learning and navigating maps when other sensory cues are absent. Similarly, encoding snapshots from a graphical model for vision [59] as the input to this sequencer might enable the learning of visuo-spatial concepts and visual routines [60], and model the bi-directional influence hippocampus has on the visual cortex [61]. We believe these ideas are promising paths for future exploration. Although beyond the scope of the current work, hippocampal replay [62] is a phenomenon that could potentially be explained using CSCGs. Our related work [63] has shown that an algorithm that rapidly memorizes and gradually generalizes is possible for learning a CSCG representation. Learning from rest time replay of sequences can help such an algorithm consolidate and generalize better. Inference time replay might be explained as the searching of trajectories to multiple goals and their vicarious evaluation.

Elucidating how cognitive maps are represented in the hippocampus, how they are acquired from a stream of experiences, and how to utilize them for prediction and planning is not only crucial to understand the inner workings of the brain, but also offers key insights into developing agents with artificial general intelligence. The CSCG model, which we introduce in this paper, provides a plausible answer to each of these questions. We expect this model to be beneficial in both neuroscience and artificial intelligence as a way to produce explicit representations that are easy to interpret and manipulate from multimodal sequential data.

## Acknowledgments

We thank Chen Sun (MIT) for helpful discussions and the use of **Fig. 5c**. We thank members of Vicarious AI for critically reading this manuscript and for insightful discussions. This work was supported by a grant from the Office of Naval Research.

## Methods

### Expectation-Maximization learning of cloned HMMs

The standard algorithm to train HMMs is the expectation-maximization (EM) algorithm [71] which in this is context is known as the Baum-Welch algorithm. cloned HMM equations require a few simple modifications with respect the HMM equations: the sparsity of the emission matrix can be exploited to only use small blocks of the transition matrix both in the E and M steps and the actions, if present, should be grouped with the next hidden state (see **Fig. 1c)**, to remove the loops and create a chain that is amenable to exact inference.

Learning a cloned HMM requires optimizing the vector of prior probabilities *π*: *π*_*u*_ = *P* (*z*_1_ = *u*) and the transition matrix *T*: *T*_*uv*_ = *P* (*z*_*n*+1_ = *v*|*z*_*n*_ = *u*). To this end, we assume the hidden states are indexed such that all the clones of the first emission appear first, all the clones of the second emission appear next, etc. Let *E* be the total number of emitted symbols. The transition matrix *T* can then be broken down into smaller submatrices *T* (*i, j*), *i, j* ∈ 1 *… E*. The submatrix *T* (*i, j*) contains the transition probabilities *P* (*z*_*n*+1_|*z*_*n*_) for *z*_*n*_ ∈ *C*(*i*) and *z*_*n*+1_ ∈ *C*(*j*) (where *C*(*i*) and *C*(*j*) respectively correspond to the hidden states (clones) of emissions *i* and *j*).

The standard Baum-Welch equations can then be expressed in a simpler form in the case of cloned HMM. The E-step recursively computes the forward and backward probabilities and then updates the posterior probabilities. The M-step updates the transition matrix via row normalization.

**E-Step**

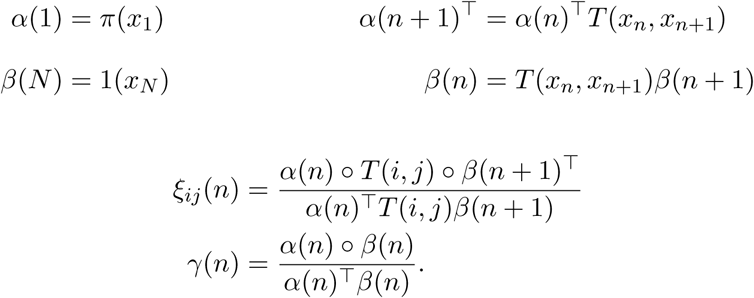

**M-Step**

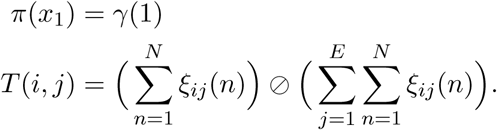

where ∘ and ∅ denote the element-wise product and division, respectively (with broadcasting where needed). All vectors are *M* × 1 column vectors, where *M* is the number of clones per emission. We use a constant number of clones per emission for simplicity here, but the number of clones can be selected independently per emission.

#### Computational savings

For a standard HMM with *H* hidden states, the computational cost for running one EM step on a sequence of length *N* is 𝒪(*H*^2^*N*) and the required memory is 𝒪(*H*^2^ + *HN*) (for the transition matrix and forward-backward messages). In contrast, a cloned HMM exploits the sparse emission matrix: with *M* clones per emission, the computational cost is 𝒪(*M* ^2^*N*) and the memory requirement is 𝒪(*H*^2^ + *MN*), in the worst case. Also, there will be additional savings for every pair of symbols that never appear consecutively in the training sequence (since the corresponding submatrix of the transition matrix does not need to be stored). Memory requirements can be improved further by using the online version of EM described in the Supplementary Materials.

Since *H* = *ME*, where *E* is the total number of symbols, an increase in alphabet size will increase the computation cost of HMMs, but will not affect the cost of cloned HMMs.

Intuitively, the computation advantage of cloned HMMs over HMMs comes from the sparse emission matrix structure. The sparsity pattern allows cloned HMMs to only consider a smaller submatrix of the transition matrix when performing training updates and inference, while HMMs must consider the entire transition matrix.

#### CSCG: Action-augmented cloned HMM

CSCGs are an extension of cloned HMMs in which an action happens at every timestep (conditional on the current hidden state) and the hidden state of the next timestep depends not only on the current hidden state, but also on the current action. The probability density function is given by eq. (3), and reproduced here for convenience

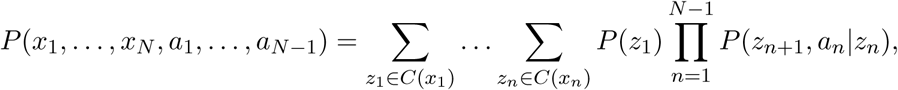

and the standard cloned HMM can be recovered by integrating out the actions. All the previous considerations about cloned HMMs apply to CSCGs and the EM equations for learning them are also very similar:

**E-Step:**

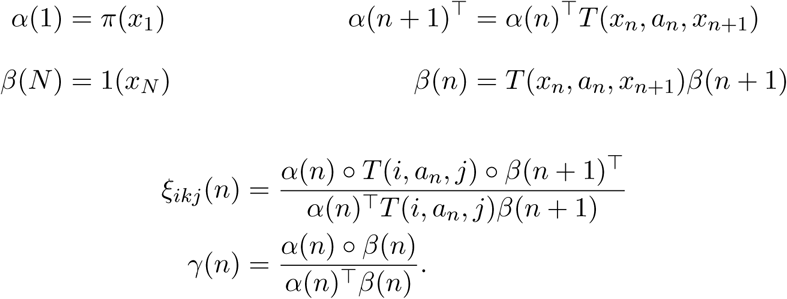

**M-Step:**

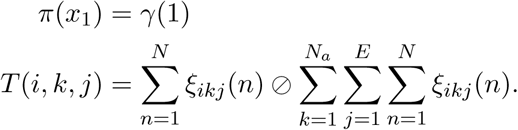

where *N*_*a*_ is the number of actions and *T* (*i, k, j*) = *P* (*z*_*n*+1_, *a*_*n*_ = *k*|*z*_*n*_) for *z*_*n*_ ∈ *C*(*i*) and *z*_*n*+1_ ∈ *C*(*j*), i.e., a portion of the action-augmented transition matrix.

#### Smoothing

We have observed that convergence can be improved by using a small pseudocount *κ*. A pseudocount is simply a small constant that is added to the accumulated counts statistic matrix 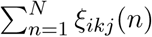 and ensures that any transition under any action has non-zero probability. This ensures that at test time the model does not have zero probability for any observations stream. When the pseudocount is only used to improve convergence, one can run EM a second time with no pseudocount, warmstarting from the result of the EM with pseudocount. To use the pseudocount, we only need to change our transition matrix update to be 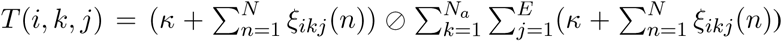. The pseudocount can be interpreted as the hyperparameter of a Laplacian prior that is set on the transition matrix, and EM as solving MAP inference for such hyperparameter. As any prior, the pseudocount has a regularization effect that helps generalization when the amount of training data is small in comparison with the capacity of the model.

It might seem at first as if adding a pseudocount would destroy the block-sparse property of the transition and therefore some of the aforementioned computational advantages of the CSCG. However, it is easy to see that the resulting transition matrix can still be expressed as the sum of a block-sparse matrix (with the same sparsity pattern as before) and a rank-1 matrix (which is not stored explicitly, but as the two vectors whose outer product produce it). By doing this, the pseudocount can be used without increasing the computational complexity or the storage requirements of any of our algorithms (EM learning, inference, etc).

#### Inference

Since the resulting model (with the action *a*_*n*_ and hidden state *z*_*n*+1_ collapsed in a single variable) forms a chain, inference on it using belief propagation (BP) is exact. When no evidence is available for a given variable, BP will simply integrate it out, so we can for instance train a model with actions and then, at test time, use it even if no actions are available. We can still ask the model which observation is the most likely in the next timestep, or even several timesteps ahead, and BP will produce the exact answer by analytically integrating over all possible past and future actions, and even over the unseen future observations when necessary.

The same model can be used to generate sequences (e.g., to generate plausible observations and actions that would correspond to wandering in a previously learned room) simply by applying ancestral sampling [70] to the conditionals that describe the model after learning (i.e., the transition and emission matrices).

A consequence of the above for spatial data is that an agent roaming the world can infer where in an environment it is located (*z*_*n*_) and then predict which actions are feasible at that location, which is useful for navigation. One can even condition on a future location to discover which set of actions can take you there, and which observations you are expected to see on the way there, see e.g. **Figs. 3b&c**. This is essentially planning as inference [34].

All of this flexible querying is performed by running a single algorithm (BP) on the same model (without retraining) and only changing the selection of which evidence is available and which probabilistic predictions are requested.

### Experimental details

#### Emergence of spatial maps from aliased sequential observations

For this experiment we collected a stream of 50000 action-observations pairs. We learned a CSCG with 20 clones (a total of 360 states) with pseudocount 2 · 10^−3^ and ran EM for 1000 iterations. This gets a result that is very close to the global minimum: when Viterbi decoded, only 48 distinct states are in use, which is the theoretical optimum on a 6 × 8 grid. Viterbi training [39] is used to refine the previous solution.

#### Transitive inference: disjoint experiences can be stitched together into a coherent whole

**Fig. 2e** showcases the CSCG’s ability to stitch together two disjoint room experiences when the rooms overlap. For this experiment, we randomly generate two square rooms of size 8 × 6 with 15 different observations each. We make both rooms share a 3 × 3 patch in their corners as shown in **Fig. 2e**.

We sample a random walk of length 10000 of action-observation pairs on each room, always avoiding to take actions that would make the random walk move outside of the room. We use 20 clones, which is enough to fully recover both rooms separately, and use a pseudocount of 10^−2^. We run EM (on both sequences simultaneously, as two independent observations of the same CSCG) for a maximum of 740 100 iterations. After EM convergence, we additionally use Viterbi decoding (with no pseudocount) to remove unused clones. The learned CSCG is visualized in **Fig. 2f**, showing that the two rooms that were experienced separately have been stitched together. Predictive performance on the stitching of the two rooms is perfect (indicating that learning succeeded) after a few observations required for the agent to locate itself. Notice that there is another patch in the first room that is identical to the merged patches, but was not merged. The model is using the sequential information to effectively identify patches that can be merged while respecting the observational data and context, and not simply looking for locally identical patches to merge.

#### Learned spatial maps form a reusable structure to explore similar environments

For this experiment, we train on a 6 × 8 room using 10000 action-observation pairs. We call this Room 1, see **Fig. 3a**. There are only 20 unique symbols in the room, some of which are repeated. The pseudocount is set to 10^−2^ and we use 20 clones (in this case, only 7 clones are strictly required to memorize the room). The regularizing pressure of the pseudocount effectively removes redundant clones. Training is done using EM for a maximum of a 100 iterations. This results in an almost perfect discovery of the underlying graph. Then we set the pseudocount to 0 and continue the training using Viterbi training [39]. This results in perfect discovery of the underlying graph with no duplicate clones. Predictions become perfect after a few initial observations required to know where in the room we are.

When we try to partially learn Room 2 with a few samples from its periphery (see **Fig. 3a**), we create a new CSCG with the transition matrix that we learned from Room 1 and keep it fixed. The emission matrix is initialized uniformly and learned using EM. The whole data for learning the emission matrix are only the 20 action-observation pairs seen in **Fig. 3a**. At that point, we fix the model and query it for a return path plan, both with and without blockers in the path. The results are displayed in **Fig. 3b** and **Fig. 3c**.

In **Fig. 3e-g** we showcase the increase in data efficiency when we transfer the learned topology to a new room with different observations. First we ignore the results from training on Room 1 and train on a new room, Room 2, from scratch following the same procedure outlined above. We train on the first *N* action-observation pairs and predict for the rest. We average (geometrically) the probability of getting the next observation right for the last 8000 samples of the 10000 available. This results in the graph in **Fig. 3e**, where N is shown in the horizontal axis. Then we repeat the same procedure, but instead of training from scratch from a random transition matrix with fixed emissions, we fix the transition matrix that we got from training in Room 1 and we learn the emission matrix, which is initialized to uniform. EM for the emission matrix converges in a few iterations. Once all nodes have been observed (when the red curve achieves 1.0), this procedure converges to perfect predictions in one or two EM iterations. This results in the graph in **Fig. 3f**, where again the horizontal axis shows *N*, the number of training action-observation pairs.

#### Representation of paths and temporal order

To learn the CSCG on the maze in **Fig. 4f**, we sample 5000 paths along each of the stochastic routes that are shown. The number in each cell indicates the observation received at that cell, and the arrows indicate possible transitions. We consider both sequences as independently generated by the model, and run EM to optimize the probability of both simultaneously. We allocate 20 clones for each other observation. By inspecting the sum-product forward messages of belief propagation at each step as the rat navigates the two routes, we can see the distributions over clones. We observe they are over disjoint subsets of the clones. To generate the paths from the CSCG shown in **Fig. 4g** (producing only paths that are consistent with the route), we sample an observation from the normalized messages from hidden state to observation during forward message passing. Finally, to extract the communities and generate the visualization in **Fig. 4g**, we run the InfoMap algorithm [69] on the graph defined by CSCG transition matrix.

In **Fig. 5**, we replicate the experiments of Sun and colleagues [21] as follows. First, we learned a CSCG with 20 clones per observation on a sequence of observations sampled from four laps around the maze shown in **Fig. 5d**. The start and end positions were unique observations. Training was terminated when perfect learning of the underlying graph was achieved. Community detection (explained below) revealed that each sensory observation was encoded by a unique clone, akin to the chunking cells found by Sun and colleagues.

#### Retrieval and Remapping

We generate random walks (random actions out of up, right, down, left) of length 10000 in each of 5 mazes. For **Fig. 6a** the mazes are 5×5 rooms where the observations are assigned to cells by a random permutation of the values 1 to 25, inclusive. For **Fig. 6b**, the structure of the mazes is shown and the observations are indicated by the color of the cells. We constructed these mazes such that have many shared observations, but each has some distinct structure that differentiates it from the others.

For each of the experiments, we learn a CSCG on these random walk sequences. After learning, we sum the forward messages of sum-product belief propagation in each maze to get a distribution over hidden states for each maze. Now on a test sequence, we can use the forward messages and these clone distributions per maze to infer the probability of being in each maze at each time step. In each of subfigure of **Fig. 6**, we shows these predictions as well as the distribution over clones over time.

Learning a CSCG in these maze environments can also enable error correction of noisy/corrupted observations. To correct errors in a corrupted observation sequence we modify the emission matrix to generate a random symbol with a small probability, thus modeling errors. Then we perform sum-product message passing on sequences with errors and find the most likely a posteriori value for each symbol. In our case, we only perform a forward pass, which provides an online estimation (based only on past data) of the MAP solution. We will use a corruption probability of 20% in our experiments, uniform over the incorrect symbols. For the 5×5 rooms, this procedure was able to correct 50 of the 55 corrupted symbols while not corrupting any of the uncorrupted symbols. For the mazes, this procedure was able to correct 46 of the 54 corrupted symbols while, again, not corrupting any of the uncorrupted symbols.

#### Community detection and hierarchical planning

In all figures, custom Python scripts were used to convert the transition matrix of the CSCG into a directed graph. This graph was then visualized using built-in functions in `python-igraph` (https://igraph.org/python/). Similarly, community detection was performed using igraph’s built-in `infomap` function.

In the hierarchical planning experiments shown in **Fig. 7**, we first generated each room by drawing a random integer between 1 and 12 with repetitions to serve as observations. The rooms were then connected via bridges (observation 13, colored black) and were tiled to form a maze as shown in **Fig. 7d**. Next, we trained a CSCG with 40 clones per observation using 1000 random restarts as described above. The learned CSCG achieved perfect prediction accuracy, suggesting perfect learning of the underlying graph. Community detection on the learned CSCG was performed using `igraph`. To form the top-level graph shown in **Fig. 7e**, we collapsed each distinct community into a single node. These communities roughly corresponded to each of the rooms in the maze. In some cases, certain rooms were partitioned into multiple communities. Next, we ran community detection on this graph to retrieve the hyper-rooms.

To compute the shortest trajectory between two locations on the maze, we first computed the shortest path in the highest-level graph using Dijkstra’s algorithm, implemented in `networkx` (https://networkx.github.io/). This returned the sequence of hyper-rooms and rooms to be visited in order to reach the goal from a start point. Next, we pruned the community graph to include only clones corresponding to these rooms and then found the shortest path in this reduced graph, which gave the exact sequence of observations from the start position to the goal (denoted by the black arrow in **Fig. 7d**. This hierarchical approach was consistently better than searching for the shortest path on the full maze itself with an average 25% fewer steps (n = 10 mazes). To determine to what extent the partitioning of the CSCG transition matrix into communities helped planning, we formed surrogate communities which no longer respected room boundaries. This resulted in a planned trajectory with an average 35% more steps than the hierarchical plan.

It is important to note that hierarchical planning is significantly more efficient. A representative example of the reduction in complexity can be given as follows. Assume that we have *V* communities, *E* inter-community edges and *M* nodes inside each community. Further, assume that the nodes inside each community are fully connected, and between any two communities, there is at most one edge. Then with hierarchical planning the complexity of running Djikstra’s algorithm is 𝒪(*E* + *V* log *V* + *N*_*t*_(*M* (*M* − 1)*/*2 + *M* log *M*)), where *N*_*t*_ is the number of top-level nodes to traverse in the second planning stage. In contrast, on the full graph, the complexity is 𝒪(*E* + *MV* log *MV* + *V* (*M* (*M* − 1)*/*2)). From these equations it is easy to see that hierarchical planning is more efficient because *N*_*t*_ ≤ *V* in all graphs.

### Adaptive and online EM variant of CSCGs

Although the sequences in the experiments of this work are not too large, we might want to be able deal with cases in which there is a very long incoming stream of observations, so long that we cannot even store it in its entirety. In order to handle this case, we can simply extend the previous EM algorithms to make them online.

The adaptive, online version of the EM algorithm in the **Methods** section is obtained by splitting the sequence in *B* batches *b* = 1 *… B* and performing EM steps on each batch successively. This allows the model to adapt to changes in the statistics if those happen over time. The statistics *ξ*_*ij*_(*n*) of batch *b* are now computed from the E-step over that batch, using the transition matrix *T* ^(*b*−1)^ from the previous batch. After processing batch *b*, we store our running statistic in *A*^(*b*)^ as:

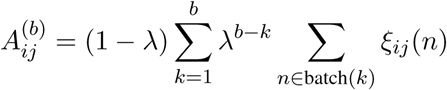

and then compute the transition matrix *T* ^(*b*)^ as

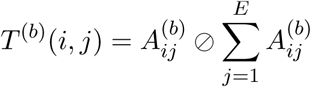

where 0 < *λ* < 1 is a memory parameter and *n* ∈ batch(*k*) refers to the time steps contained in batch *k*. For *λ* → 1, *T* ^(*b*)^(*i, j*) coincides with the transition matrix from the **Methods** section. For smaller values of *λ*, the expected counts are weighed using an exponential window,^1^ thus giving more weight to the more recent counts.

To learn from arbitrarily long sequences, we consider an online formulation and express 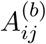 recursively:

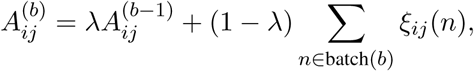

so that the expected counts of the last observed batch are incorporated into the running statistics.

## Data and code availability

All python scripts used to simulate and analyze the model will be deposited on GitHub.

## Supplementary Results

**Supplementary Fig. 1:**
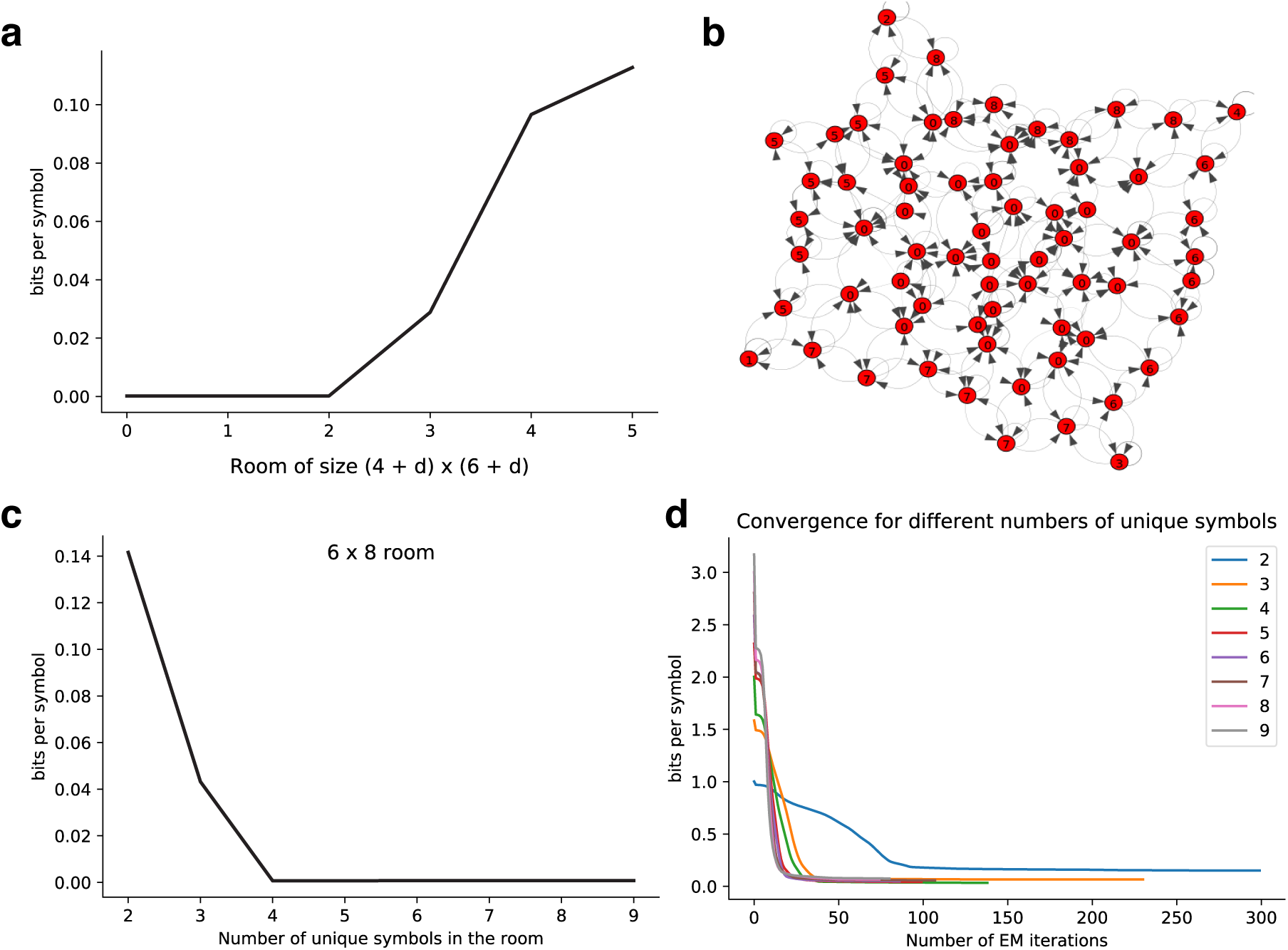
The effect of varying room size and the number of unique observations in each room. **(a)** Larger empty rooms result in increased bps. **(b)** Imperfect graph learned from a large empty room. **(c)** Fewer unique observations result in increased bps. **(d)** Fewer unique observations make learning harder.

### Learning spatial representations

In order to learn the spatial representation of a room with a CSCG, we let an agent roam in a room and receive a stream of local visual cues paired with the executed actions. Note that there are two factors that complicate learning: on the one hand, the visual cues do not need to be unique to each location in the room. In fact, in an empty room, every location in the room away from the walls and the corners looks the same (see Fig. 2c). On the other hand, even though the agent’s proprioception lets it know the identifier of the executed action, there is no meaning associated to it. I.e., every time that the agent moves west, it knows it is doing the same thing, but it does not have any prior knowledge of what that thing is.

In the case of empty rooms, most of the observations received by the agent are the same (the empty observation, as it wanders through empty space) and it is hard for the agent to locate itself in the room, since only the walls and corners provide context. We have experimented with CSCGs learning in empty rooms of different sizes. We use 50000 steps^2^ in a room of size (4 + *d*) × (6 + *d*), where *d* is a parameter controlling the room size. For rooms of size 6 × 8 and below, EM learning recovers exactly the structure of the room. For larger sizes, it starts to make some mistakes in its understanding of the room, slightly decreasing its predictive ability as the room grows, see **Supplementary Fig. 1a. Supplementary Fig. 1b** shows the learned transition matrix in graph form for a room of size 9 × 11. The graph looks almost perfect, but if we follow the path between observations ‘1’ and ‘3’, we should traverse seven observations of type ‘7’, whereas there are only six. The CSCG has merged two physical locations in the room (that have a large neighborhood of identical sensory cues) into the same perceived location.

Learning the structure of a room would be trivial if each observation was unique to a single room location. In that case, a CSCG with only one clone would learn the correct solution in one EM step. We experiment with different numbers of unique symbols randomly placed in a room. **Supplementary Fig. 1c** shows that in a room of size 6 × 8 (depicted in **Fig. 2a**) the performance degrades as the number of unique symbols decreases, with recovery being exact only when the number of unique symbols is 4 or more. The number of EM iterations required for convergence^3^ is also affected, as shown in **Supplementary Fig. 1d**.

### Successor representation of the CSCG transition matrix

The CSCG model contains all the information about the sequence generating process, so it can be combined with an external policy to yield the successor representation associated to that policy. The successor representation[9, 82] loses precise temporal information and, as a result, contains strictly less information than the CSCG. Additionally, unlike the CSCG, the successor representation assumes full observability of the state, so it cannot be derived from partial or aliased observations.

**Supplementary Fig. 2:**
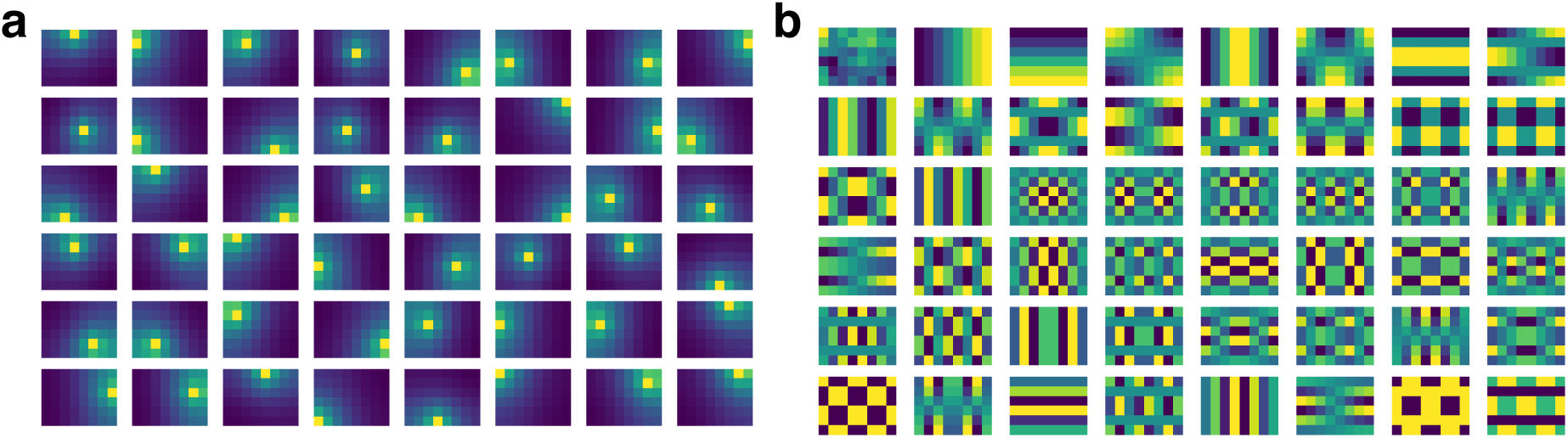
Successor representation and eigenvectors derived from the transition matrix of a CSCG. The CSCG was trained with data collected from a random walk in a rectangular room. **(a)** Successor Representations in 6×8 room. **(b)** Eigenvectors of Successor Representation.

**Supplementary Fig. 3:**
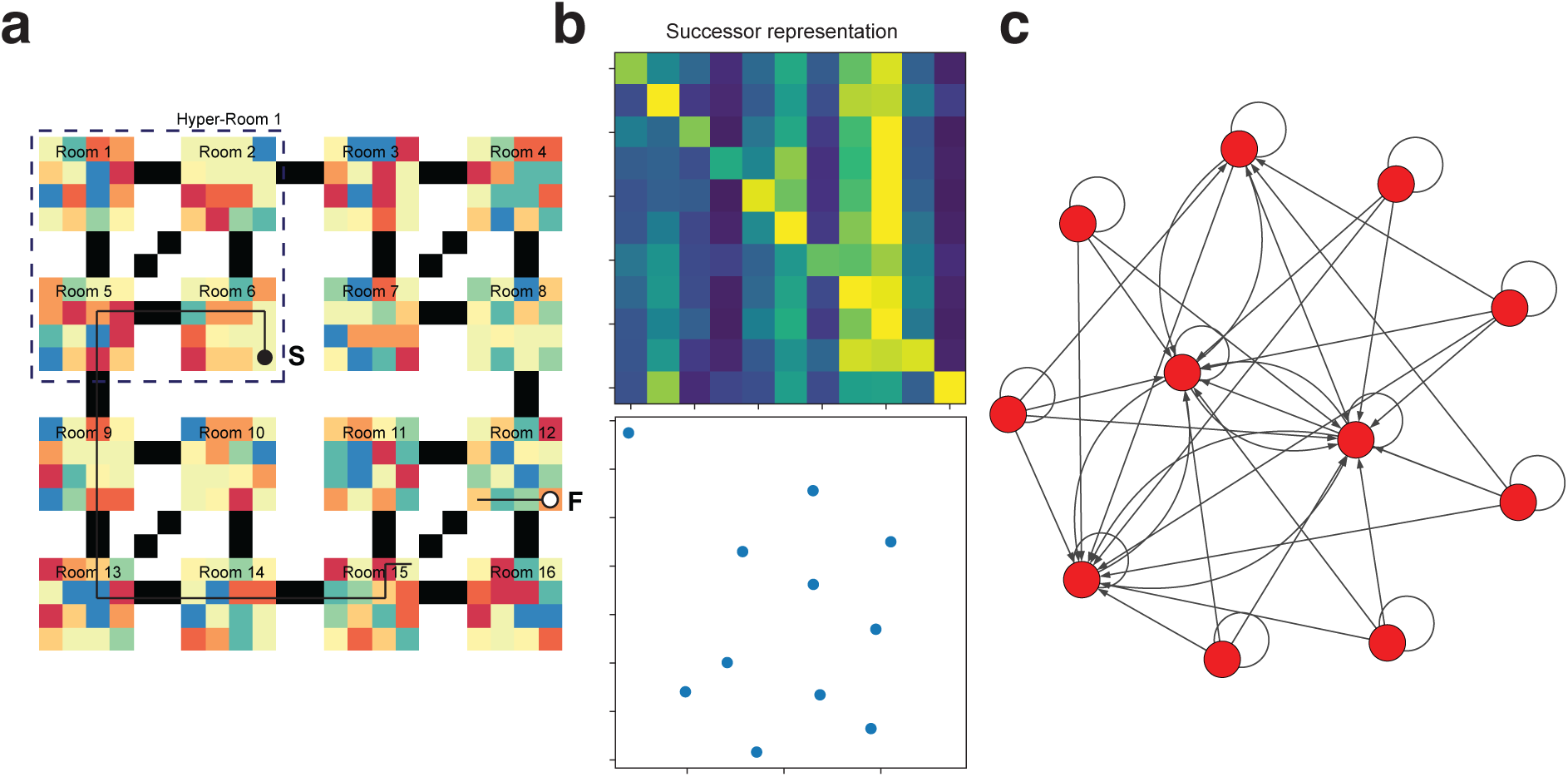
Successor representation for the hierarchical maze experiment. **(a)** Same maze as shown in **Fig. 7d. (b)** Successor Representation matrix and MDS. **(c)** Graph derived from Successor Representation Matrix.

We take the CSCG learned from the aliased 6×8 room in **Fig. 2a** and generate the SR from the CSCG transition matrix. Then we identify which clones correspond to which spatial locations by observing which clones activate in each location during inference. In **Supplementary Fig. 2a** we visualize the SR for each hidden state. We also compute the eigenvectors of the matrix containing the SR of each state. We visualize these eigenvectors in **Supplementary Fig. 2b** and we also observe various grid patters of different scales, similarly to [9].

**Supplementary Video 1** The left panel shows the physical location of the agent and the local visual cue (color) available to it, whereas the right panel shows the inferred position in the agent’s cognitive map (which has been learned from data). The agent only observes the current color (and not even its own actions). There are two patches (marked in black) that have identical colors, so at the beginning of exploration, the agent’s belief in the cognitive map (right) is split between the two possible realities. As soon as the agent exits the duplicated patch, it can figure out its precise location and track it properly from that point on, as shown by the lack of ambiguity in the cognitive map when the agent returns to the repeated patch.

**Supplementary Video 2** Inferred cognitive map over EM iterations. The CSCG transition matrix is updated after each EM iteration, and the current state of the model is displayed as a cognitive map. To do this, the training data is decoded as a sequence of clones using Viterbi, and the resulting clone transitions are represented in a graph. The layout of the graph is obtained automatically using `python-igraph`.

## Footnotes

1 Normalization of the exponential window is unnecessary, since it will cancel when computing *T* ^(*b*)^(*i, j*).

2 Other parameters for this experiment are the number of clones used (70, which is theoretically enough to exactly recover even the largest room), a pseudocount of 2 · 10^−3^ used in the EM procedure and a maximum number of EM iterations of 1000.

3 The parameters for this experiment are: 10000 recorded steps, a CSCG with 30 clones (enough for exact recovery for all the numbers of unique symbols tested), a pseudocount of 10^−2^ in the EM procedure and a maximum number of EM iterations of 1000.

## References

1. Redish, A. D. Vicarious trial and error. Nature Reviews Neuroscience 17, 147 (2016).

2. Tolman, E. C. Cognitive maps in rats and men. Psychological review 55, 189 (1948).

3. Epstein, R. A., Patai, E. Z., Julian, J. B. & Spiers, H. J. The cognitive map in humans: spatial navigation and beyond. Nature neuroscience 20, 1504 (2017).

4. Buzsáki, G. & Llinás, R. Space and time in the brain. Science 358, 482–485 (2017).

5. Niv, Y. Learning task-state representations. Nature neuroscience 22, 1544–1553 (2019).

6. Schapiro, A. C., Turk-Browne, N. B., Norman, K. A. & Botvinick, M. M. Statistical learning of temporal community structure in the hippocampus. Hippocampus 26, 3–8 (2016).

7. Eichenbaum, H., Dudchenko, P., Wood, E., Shapiro, M. & Tanila, H. The hippocampus, memory, and place cells: is it spatial memory or a memory space? Neuron 23, 209–226 (1999).

8. Moser, E. I., Kropff, E. & Moser, M.-B. Place cells, grid cells, and the brain’s spatial representation system. Annu. Rev. Neurosci. 31, 69–89 (2008).

9. Stachenfeld, K. L., Botvinick, M. M. & Gershman, S. J. The hippocampus as a predictive map. Nature neuroscience 20, 1643 (2017).

10. Dayan, P. Improving Generalization for Temporal Difference Learning: The Successor Representation. Neural Computation 5, 613–624. eprint: https://doi.org/10.1162/neco.1993.5.4.613. https://doi.org/10.1162/neco.1993.5.4.613 (1993).

11. Piray, P. & Daw, N. D. A common model explaining flexible decision making, grid fields and cognitive control. bioRxiv. eprint: https://www.biorxiv.org/content/early/2019/12/10/856849.full.pdf. https://www.biorxiv.org/content/early/2019/12/10/856849 (2019).

12. Whittington, J., Muller, T., Mark, S., Barry, C. & Behrens, T. in Advances in Neural Information Processing Systems 31 (eds Bengio, S. et al.) 8484–8495 (Curran Associates, Inc., 2018). http://papers.nips.cc/paper/8068-generalisation-of-structural-knowledge-in-the-hippocampal-entorhinal-system.pdf.

13. Colgin, L. L., Moser, E. I. & Moser, M.-B. Understanding memory through hippocampal remapping. Trends in neurosciences 31, 469–477 (2008).

14. Aronov, D., Nevers, R. & Tank, D. W. Mapping of a non-spatial dimension by the hippocampal– entorhinal circuit. Nature 543, 719 (2017).

15. Wood, E. R., Dudchenko, P. A., Robitsek, R. J. & Eichenbaum, H. Hippocampal neurons encode information about different types of memory episodes occurring in the same location. Neuron 27, 623–633 (2000).

16. Frank, L. M., Brown, E. N. & Wilson, M. Trajectory encoding in the hippocampus and entorhinal cortex. Neuron 27, 169–178 (2000).

17. Duvelle, É. et al. Insensitivity of place cells to the value of spatial goals in a two-choice flexible navigation task. Journal of Neuroscience 39, 2522–2541 (2019).

18. Gauthier, J. L. & Tank, D. W. A dedicated population for reward coding in the hippocampus. Neuron 99, 179–193 (2018).

19. Wills, T. J., Lever, C., Cacucci, F., Burgess, N. & O’Keefe, J. Attractor dynamics in the hippocampal representation of the local environment. Science 308, 873–876 (2005).

20. Grieves, R. M., Wood, E. R. & Dudchenko, P. A. Place cells on a maze encode routes rather than destinations. Elife 5, e15986 (2016).

21. Sun, C., Yang, W., Martin, J. & Tonegawa, S. CA1 pyramidal cells organize an episode by segmented and ordered events (2019).

22. Cormack, G. V. & Horspool, R. N. S. Data compression using dynamic Markov modelling. The Computer Journal 30, 541–550 (1987).

23. Xu, J., Wickramarathne, T. L. & Chawla, N. V. Representing higher-order dependencies in networks. Science advances 2, e1600028 (2016).

24. Hawkins, J., George, D. & Niemasik, J. Sequence memory for prediction, inference and behaviour. Philosophical Transactions of the Royal Society B: Biological Sciences 364, 1203–1209. ISSN: 14712970. http://www.ncbi.nlm.nih.gov/pubmed/19528001%20http://www.pubmedcentral.nih.gov/articlerender.fcgi?artid=PMC2666719 (May 2009).

25. Cui, Y., Ahmad, S. & Hawkins, J. Continuous online sequence learning with an unsupervised neural network model. Neural Computation 28, 2474–2504. ISSN: 1530888X. 1512.05463. http://arxiv.org/abs/1512.05463 (Dec. 2016).

26. Dedieu, A. et al. Learning higher-order sequential structure with cloned HMMs. 1905.00507. http://arxiv.org/abs/1905.00507 (May 2019).

27. Sharan, V., Kakade, S. M., Liang, P. S. & Valiant, G. Learning overcomplete hmms in Advances in Neural Information Processing Systems (2017), 940–949.

28. Palacios, E. R., Razi, A., Parr, T., Kirchhoff, M. & Friston, K. On Markov blankets and hierarchical self-organisation. Journal of theoretical biology 486, 110089 (2020).

29. Manning, C. D., Raghavan, P. & Schütze, H. Introduction to information retrieval (Cambridge university press, 2008).

30. Rao, R. P. Bayesian computation in recurrent neural circuits. Neural computation 16, 1–38 (2004).

31. George, D. & Hawkins, J. Towards a mathematical theory of cortical micro-circuits. PLoS computational biology 5 (2009).

32. Nessler, B., Pfeiffer, M., Buesing, L. & Maass, W. Bayesian computation emerges in generic cortical microcircuits through spike-timing-dependent plasticity. PLoS computational biology 9 (2013).

33. Whittington, J. C. et al. The Tolman-Eichenbaum Machine: Unifying space and relational memory through generalisation in the hippocampal formation. bioRxiv, 770495 (2019).

34. Attias, H. Planning by probabilistic inference. in AISTATS (2003).

35. Morton, N. W., Sherrill, K. R. & Preston, A. R. Memory integration constructs maps of space, time, and concepts. Current opinion in behavioral sciences 17, 161–168 (2017).

36. Behrens, T. E. et al. What is a cognitive map? Organizing knowledge for flexible behavior. Neuron 100, 490–509 (2018).

37. Baraduc, P., Duhamel, J.-R. & Wirth, S. Schema cells in the macaque hippocampus. Science 363, 635–639 (2019).

38. Blum, K. I. & Abbott, L. A model of spatial map formation in the hippocampus of the rat. Neural computation 8, 85–93 (1996).

39. Jelinek, F. Continuous speech recognition by statistical methods. Proceedings of the IEEE 64, 532–556 (1976).

40. Ginther, M. R., Walsh, D. F. & Ramus, S. J. Hippocampal neurons encode different episodes in an overlapping sequence of odors task. Journal of Neuroscience 31, 2706–2711 (2011).

41. Alme, C. B. et al. Place cells in the hippocampus: eleven maps for eleven rooms. Proceedings of the National Academy of Sciences 111, 18428–18435 (2014).

42. Lever, C., Wills, T., Cacucci, F., Burgess, N. & O’Keefe, J. Long-term plasticity in hippocampal place-cell representation of environmental geometry. Nature 416, 90–94 (2002).

43. Latuske, P., Kornienko, O., Kohler, L. & Allen, K. Hippocampal remapping and its entorhinal origin. Frontiers in behavioral neuroscience 11, 253 (2018).

44. Sosa, M., Gillespie, A. K. & Frank, L. M. in Behavioral Neuroscience of Learning and Memory 43–100 (Springer, 2016).

45. Leutgeb, J. K. et al. Progressive transformation of hippocampal neuronal representations in “morphed” environments. Neuron 48, 345–358 (2005).

46. Balaguer, J., Spiers, H., Hassabis, D. & Summerfield, C. Neural mechanisms of hierarchical planning in a virtual subway network. Neuron 90, 893–903 (2016).

47. Tomov, M. S., Yagati, S., Kumar, A., Yang, W. & Gershman, S. J. Discovery of Hierarchical Representations for Efficient Planning. bioRxiv. eprint: https://www.biorxiv.org/content/early/2019/03/28/499418.full.pdf. https://www.biorxiv.org/content/early/2019/03/28/499418 (2019).

48. Bohlin, L., Edler, D., Lancichinetti, A. & Rosvall, M. in Measuring scholarly impact 3–34 (Springer, 2014).

49. Zhang, P. & Moore, C. Scalable detection of statistically significant communities and hierarchies, using message passing for modularity. Proceedings of the National Academy of Sciences 111, 18144–18149 (2014).

50. Buzsáki, G. & Tingley, D. Space and time: The hippocampus as a sequence generator. Trends in cognitive sciences 22, 853–869 (2018).

51. Whittington, J., Muller, T., Mark, S., Barry, C. & Behrens, T. Generalisation of structural knowledge in the hippocampal-entorhinal system in Advances in neural information processing systems (2018), 8484–8495.

52. Momennejad, I. et al. The successor representation in human reinforcement learning. Nature Human Behaviour 1, 680 (2017).

53. Momennejad, I. & Howard, M. W. Predicting the future with multi-scale successor representations. BioRxiv, 449470 (2018).

54. Kansky, K. et al. Schema networks: Zero-shot transfer with a generative causal model of intuitive physics in Proceedings of the 34th International Conference on Machine Learning-Volume 70 (2017), 1809–1818.

55. Holmes, M. P. et al. Schema learning: Experience-based construction of predictive action models in Advances in Neural Information Processing Systems (2005), 585–592.

56. Summerfield, C., Luyckx, F. & Sheahan, H. Structure learning and the posterior parietal cortex. Progress in neurobiology, 101717 (2019).

57. Wilson, R. C., Takahashi, Y. K., Schoenbaum, G. & Niv, Y. Orbitofrontal cortex as a cognitive map of task space. Neuron 81, 267–279 (2014).

58. Pezzulo, G., Cartoni, E., Rigoli, F., Pio-Lopez, L. & Friston, K. Active Inference, epistemic value, and vicarious trial and error. Learning & Memory 23, 322–338 (2016).

59. George, D. et al. A generative vision model that trains with high data efficiency and breaks text-based CAPTCHAs. Science 358, eaag2612 (2017).

60. Lázaro-Gredilla, M., Lin, D., Guntupalli, J. S. & George, D. Beyond imitation: Zero-shot task transfer on robots by learning concepts as cognitive programs. arXiv preprint 1812.02788 (2018).

61. Saleem, A. B., Ayaz, A., Jeffery, K. J., Harris, K. D. & Carandini, M. Integration of visual motion and locomotion in mouse visual cortex. Nature neuroscience 16, 1864 (2013).

62. Skaggs, W. E. & McNaughton, B. L. Replay of neuronal firing sequences in rat hippocampus during sleep following spatial experience. Science 271, 1870–1873 (1996).

63. Rikhye, R. V., Guntupalli, J. S., Gothoskar, N., Lázaro-Gredilla, M. & George, D. V. Memorize-Generalize: An online algorithm for learning higher-order sequential structure with cloned Hidden Markov Models. bioRxiv, 764456 (2019).

64. Friston, K. J., Parr, T. & de Vries, B. The graphical brain: belief propagation and active inference. Network Neuroscience 1, 381–414 (2017).

65. Osaba, E., Del Ser, J., Camacho, D., Bilbao, M. N. & Yang, X.-S. Community detection in networks using bio-inspired optimization: Latest developments, new results and perspectives with a selection of recent meta-heuristics. Applied Soft Computing 87, 106010 (2020).

66. Sanders, H., Wilson, M. A. & Gershman, S. J. Hippocampal Remapping as Hidden State Inference tech. rep. (Center for Brains, Minds and Machines (CBMM), bioRxiv, 2019).

67. Wilson, M. A. & McNaughton, B. L. Dynamics of the hippocampal ensemble code for space. Science 261, 1055–1058 (1993).

68. Thompson, L. & Best, P. Place cells and silent cells in the hippocampus of freely-behaving rats. Journal of Neuroscience 9, 2382–2390 (1989).

69. Rosvall, M. & Bergstrom, C. T. Maps of random walks on complex networks reveal community structure. Proceedings of the National Academy of Sciences 105, 1118–1123 (2008).

70. Bishop, C. M. Pattern recognition and machine learning (springer, 2006).

71. Wu, C. J. et al. On the convergence properties of the EM algorithm. The Annals of statistics 11, 95–103 (1983).

72. Ranganath, C. & Hsieh, L.-T. The hippocampus: a special place for time. Annals of the New York Academy of Sciences 1369, 93–110 (2016).

73. Ekstrom, A. D. & Ranganath, C. Space, time, and episodic memory: The hippocampus is all over the cognitive map. Hippocampus 28, 680–687 (2018).

74. Agster, K. L., Fortin, N. J. & Eichenbaum, H. The hippocampus and disambiguation of overlapping sequences. Journal of Neuroscience 22, 5760–5768 (2002).

75. Howard, M. W. & Eichenbaum, H. The hippocampus, time, and memory across scales. Journal of Experimental Psychology: General 142, 1211 (2013).

76. Tolman, E. C. Prediction of vicarious trial and error by means of the schematic sowbug. Psychological Review 46, 318 (1939).

77. Dordek, Y., Soudry, D., Meir, R. & Derdikman, D. Extracting grid cell characteristics from place cell inputs using non-negative principal component analysis. Elife 5, e10094 (2016).

78. Quiroga, R. Q., Reddy, L., Kreiman, G., Koch, C. & Fried, I. Invariant visual representation by single neurons in the human brain. Nature 435, 1102 (2005).

79. Tolman, E. C. There is more than one kind of learning. Psychological review 56, 144 (1949).

80. Keefe, J. O. & Nadel, L. The hippocampus as a cognitive map (Clarendon Press, 1978).

81. Ferbinteanu, J. & Shapiro, M. L. Prospective and retrospective memory coding in the hippocampus. Neuron 40, 1227–1239 (2003).

82. Gershman, S. J. The successor representation: its computational logic and neural substrates. Journal of Neuroscience 38, 7193–7200 (2018).

83. Mok, R. M. & Love, B. C. A non-spatial account of place and grid cells based on clustering models of concept learning. Nature communications 10, 1–9 (2019).

